# Brain-derived neurotrophic factor amplifies neuron-intrinsic programs to enhance axonal regeneration in human motor neurons

**DOI:** 10.1101/2023.11.06.565775

**Authors:** Jose Norberto S. Vargas, Anna-Leigh Brown, Kai Sun, Cathleen Hagemann, Bethany Geary, David Villarroel-Campos, Sam Bryce-Smith, Matteo Zanovello, Madeline Lombardo, Stan Majewski, Andrew Tosolini, Maria Secrier, Matthew J. Keuss, Andrea Serio, James N. Sleigh, Pietro Fratta, Giampietro Schiavo

## Abstract

The cell-intrinsic capacity of neurons to regenerate axons requires widespread coordination of the transcriptome, activation of multiple kinases, and reorganization of the cytoskeleton. Axonal repair is also influenced by extrinsic activating factors, such as neurotrophins. Here, we reveal that brain-derived neurotrophic factor (BDNF) amplifies multiple neuron-intrinsic programs to foster axonal regeneration in human motor neurons. Through metabolic RNA sequencing and phosphoproteomic profiling, we elucidate BDNF signalling and its role in axonal regeneration. We discover that BDNF controls RNA stability and transcriptional programs that converge with regeneration-associated gene (RAG) sets. We further unveil that BDNF governs the phosphorylation of multiple proteins essential for cytoskeletal dynamics, a major determinant of effective nerve regeneration. Using compartmentalized neuronal cultures, we demonstrate that the regeneration driven by BDNF depends on the axon-specific activation of ERK/RSK/S6K kinase pathway. We propose a model in which BDNF augments neuron-intrinsic pathways to drive axonal regeneration in human motor neurons.

**Teaser:** BDNF aids nerve repair by fine-tuning the metabolism of RNA and by changing the building blocks of the nerve cell cytoskeleton.

## Introduction

Nerve regeneration involves neuron-intrinsic pathways, as well as extrinsic activating signals, such as those provided by neurotrophins (*1–3*). The neurotrophin brain-derived neurotrophic factor (BDNF) is essential for motor neuron axon development and regeneration, as well as neuromuscular junction (NMJ) maintenance (*3*). At the NMJ, BDNF is released by the skeletal muscle and Schwann cells. BDNF then binds its cognate receptor, tropomyosin receptor kinase B (TrkB), on motor neurons, resulting in local and long-range pro-survival signalling important for nerve repair (*3*, *4*). Conditional deletion of TrkB in neurons severely impedes peripheral nerve regeneration (*5*) and lentiviral transduction of TrkB enhances the regeneration of corticospinal axons (*6*), demonstrating the crucial role of BDNF in axon regeneration. Furthermore, dysfunctional neurotrophic support is an underpinning pathomechanism for motor neuron diseases and peripheral neuropathies (*7*), further underscoring its importance for axonal maintenance.

BDNF is known to promote pro-survival gene expression through the activation of several transcription factors. Effective gene expression requires the synchronisation of RNA transcription, stability, and decay processes. Whether neurotrophins regulate not just transcription, but also control other aspects of RNA dynamics, such as stability and degradation, is currently unknown. Indeed, the regulation of the axonal and dendritic transcriptome, which is crucial for their function, can be shaped by the stability of RNAs (*8*). Advances in RNA sequencing methodologies, such as metabolic labelling approaches, allow for the direct observation of rates of RNA transcription and/or degradation (*9*). Still, to the best of our knowledge, these approaches have not been utilised to assess RNA metabolism governed by neurotrophins. Importantly, the transcriptional outputs of BDNF in human motor neurons relevant for axonal regeneration are not fully elucidated.

Another crucial gap in our understanding of BDNF/TrkB signal transduction pathway is the lack of deep profiling of the phosphorylation landscape induced by BDNF activation, particularly in human motor neurons. It is well-established that several kinases are activated by BDNF (*10*). This activation plays a key role in modulating the morphogenesis of several neuronal sub-compartments, such as growth cones, synapses, dendrites and axons, all of which depend on cytoskeletal remodelling (*11*, *12*). Indeed, axonal regeneration requires the dynamic restructuring of the cytoskeletal architecture (*13*). The organisation and function of the latter is, in part, dictated by microtubule-associated proteins (MAPs) and actin-binding proteins (*14*). Importantly, the phosphorylation of MAPs and actin-binding proteins regulates their binding to microtubules and motor proteins, thereby modulating cytoskeletal-related processes in a spatiotemporal manner (*15*). Thus, while we know that multiple kinases are activated by BDNF, we do not yet fully understand the specific phosphorylation substrates that enable BDNF to reorganise the cytoskeleton *en masse* to control neuronal architecture and axonal regeneration.

In this study, we unravel the RNA and proteomic programs controlled by BDNF through a combination of metabolic sequencing and phosphoproteomic analysis. We find a striking convergence of neuron-intrinsic regeneration pathways with those that are controlled by BDNF. We show that BDNF promotes global hypertranscription, which is a hallmark of regenerating tissues. Further, we show that BDNF governs the posttranscriptional tuning of steady-state RNA levels by altering RNA stability. Importantly, we uncover that the genetic regulatory networks controlled by BDNF overlap with those involved in promoting axonal outgrowth and repair. We further reveal that BDNF kinase signalling reshapes the phosphorylation pattern of cytoskeletal proteins, particularly of structural MAPs and actin-binding proteins. Through phosphoproteomic profiling of BDNF signalling and axonal regeneration assays using compartmented axon-specific kinase inhibition, we discovered that the specific activation of ERK-RSK-S6K pathway in axons is required for BDNF to enhance axonal regeneration in human motor neurons.

Thus, we report a comprehensive characterisation of the transcriptomic and phosphoproteomic programs controlled by BDNF signalling. These findings provide novel molecular insights as to how an extrinsic growth-promoting signal, such as BDNF, drives neuron-intrinsic regeneration programs in human motor neurons.

## Results

### Assessing RNA dynamics with SLAM-seq in human motor neurons

To study the role of BDNF in human motor neurons, we used the genetically-engineered human iPSC-derived lower motor neurons (i^3^ LMNs) expressing *NGN2*, *ISL1*, and *LHX3* under a doxycycline-inducible promoter (**Fig. S1A-D**; (*16*). Post-differentiation, i^3^ LMNs express ChAT and Hb9, which are two canonical motor neuron markers (**Fig. S1C-D**), and robustly respond to BDNF through the activation of well-known downstream kinases MAPK-ERK1/2 (referred as ERK from here), AKT, and S6K (**Fig. S1E-F**).

To dissect the transcription kinetics driven by BDNF in i^3^ LMNs, we performed SLAM-seq (*17*) 1, 2, and 6 hours of post-treatment with BDNF (**Fig. 1A**). This approach relies on the incorporation of 4-thiouridine (4sU) into nascently transcribed RNA. RNA is extracted and treated with iodoacetamide, leading to thiol-alkylation of 4sU. When RNA is mapped to the genome, this results in T>C conversions in the nascently transcribed RNA (*17*, *18*). SLAM-seq thus provides a direct readout of RNA transcription along with differentially expressed genes (DEGs) in response to BDNF. From these measurements, RNA degradation/destabilisation can be inferred by comparing ratios of newly transcribed RNA and total RNA levels, thus allowing us to investigate the effects of BDNF signalling on RNA metabolism.

**Figure 1.**
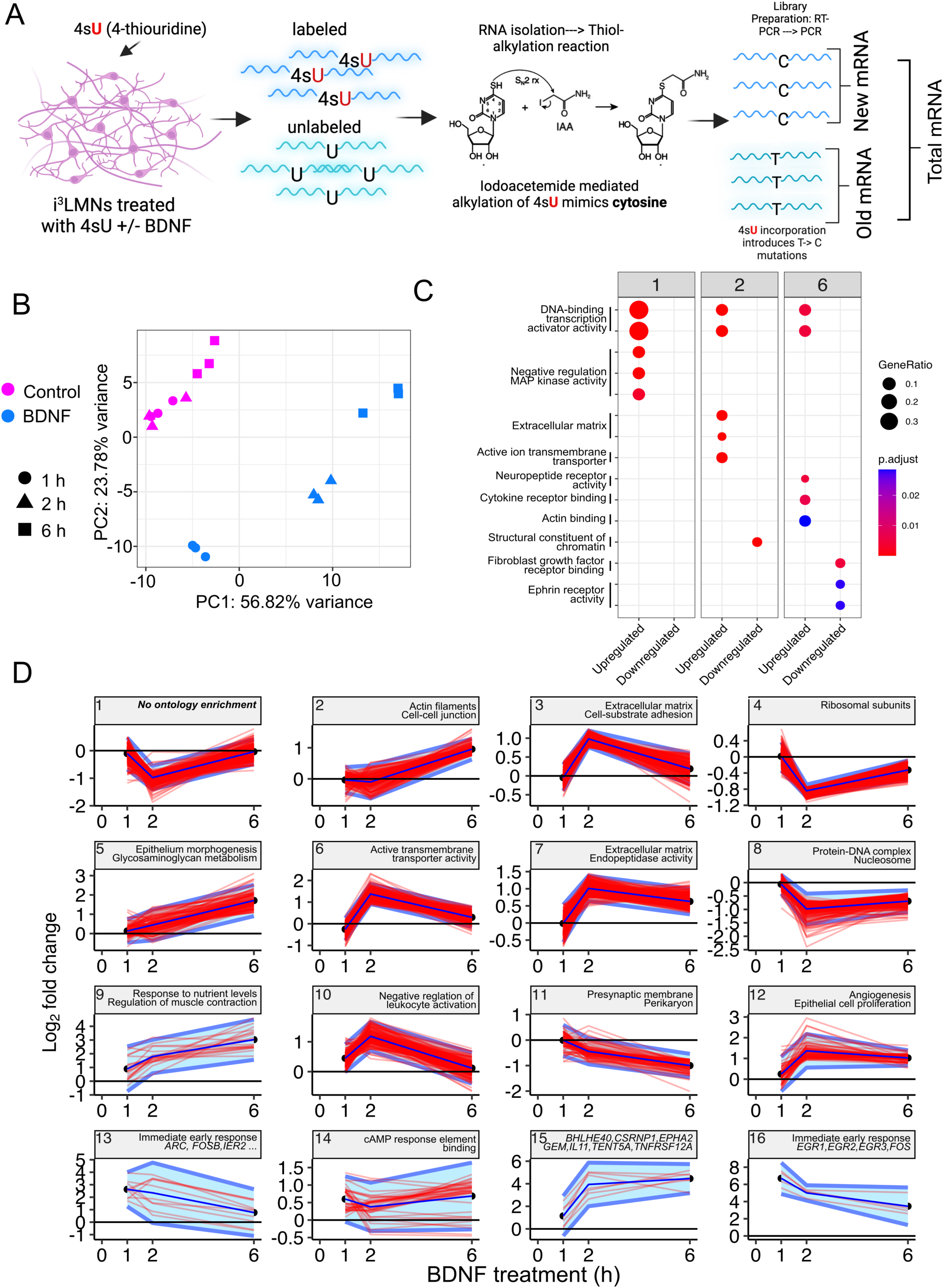
BDNF-driven transcription in human motor neurons regulates time-resolved functional networks. A) Schematic of SLAM-sequencing pipeline. Neurons were treated with 100 µM 4sU uridine analogue which is integrated into newly transcribed RNA. After RNA extraction, RNA was incubated with iodoacetamide, which promotes the thiol-alkylation of 4sU. Before standard RNA-sequencing, RNA is reverse-transcribed to cDNA; during this process the thiol-alkylated 4sU will base pair with guanine rather than adenine, thus mimicking cytosine. When the RNA-sequencing is mapped back to the genome, there will be characteristic T > C mismatches in the reads produced during the labelling period which can be used to infer the ratio of new to total RNA. N=3 biological replicates per condition and labelling time. B) Total RNA principal component analysis showing separation of control (pink) and BDNF treated (blue) lower motor neurons by both treatment condition and time. Close clustering of control samples along PC2 suggests that 4sU treatment had minimal effect on total gene abundance C) Gene-ontology analyses for up- and downregulated genes across 1, 2, 6 h BDNF treatment collapsed for semantic similarity. Differentially expressed genes on total RNA called with an adjusted p-value < 0.1 and the abs(log_2_ fold change) > 0.75 estimated by DESeq2(*62*). D) Differential gene expression hierarchically clustered by log_2_ fold change across BDNF treatment showing mean and individual gene cluster expression with cluster GO terms/gene names. Individual gene expression as red lines, mean expression of cluster dark blue lines, and twice width cluster standard deviation as light blue boundaries. Genes which were never called as differentially expressed at any of the three timepoints and with a baseMean < 15 were removed prior to clustering.

### BDNF controls functionally distinct and time-resolved transcriptional networks

First, we assessed DEGs upon BDNF treatment in human motor neurons. Principal component analysis (PCA) on total RNA revealed a clear separation between BDNF-treated and control samples (**Fig. 1B**). Immediate early response genes, reflecting a rapid but transient response to BDNF, are the main drivers of variance in condition- and time-correlated principal components (**Fig. S2A**). We detected the high expression of *EGR1*, *FOS*, *JUNB*, and *ARC,* which are known early-responsive genes to BDNF signalling, confirming the validity of both the treatment and sequencing (**Fig. S2B**). Overall, we found that the number of BDNF-dependent DEGs increased from 1 (58 genes) to 2 hours (2,131 genes), followed by a reduction at 6 hours (604 genes) (**Fig. S2B-D**; **Table S1**), suggesting that gene expression driven by extrinsic BDNF exhibits autoregulation at longer timepoints.

To functionally map the transcriptional output of BDNF, we next performed gene-ontology (GO) enrichment for DEGs across labelling times (**Fig. 1C**). DNA-binding transcription activators are significantly enriched across all timepoints, consistent with the role of BDNF in transcriptional control. Negative regulators of the MAPK pathway *DUSP1*, *DUSP5*, and *DUSP6* are transiently upregulated at 1 hour (**Fig. 1C; Fig. S2B-D**), potentially indicating a negative feedback transcriptional response to regulate sustained ERK signalling. DEGs related to extracellular matrix (ECM) and neuronal cytoskeletal components, including actin-binding factors, are enriched at later time points (**Fig. 1C; Table S2**). At 2 hours, chromatin structural genes were downregulated, perhaps reflecting BDNF’s effect on chromatin accessibility to induce gene expression (*19*). Genes encoding members of the EPHR receptor tyrosine kinase family (*EPHA4*/*EPHA3*), which are mainly axon-repulsive, are downregulated by 6 hours, suggesting that prolonged BDNF treatment can negatively regulate axonal repulsion (**Fig. 1C**; **Table S2**). Intriguingly, reduction of *EPHA4* increases peripheral nervous system regeneration (*20*) and lower *EPHA4* expression has been shown to protect against axonal degeneration in both animal models and ALS patients during disease progression (*21*).

BDNF/TrkB signalling is tightly regulated, with its effects on neuronal morphology and differentiation varying based on treatment duration and BDNF concentration (*12*). Furthermore, BDNF-driven gene expression in mouse and human cortical neurons has been shown to exhibit distinct early and late transcriptional responses (*19*). To explore this phenomenon in motor neurons, we analysed temporal and functional RNA expression downstream of BDNF signalling at different time points using hierarchical clustering and identified 16 distinct expression clusters (**Fig. 1D; Table S2**). Immediate early response genes (clusters 13 and 16) showed a rapid upregulation, returning to baseline by 6 hours, indicating the validity of this approach. Genes related to neuronal morphology, such as actin filaments (cluster 2) and ECM (clusters 3 and 7), had delayed responses, peaking after 1 hour and increasing over time. Clusters involving ribosomal subunits (cluster 4) and DNA-binding components (cluster 8) were downregulated at 2 hours but began to rebound by 6 hours. A small cluster (cluster 15) exhibited sustained upregulation (**Fig. 1D; Table S2**). These data show highly specific clusters of genes that respond in a coordinated manner to BDNF. This indicates that the temporal activation of TrkB regulates specific cellular processes through functionally-defined transcriptional responses. Of interest, we observed gene clusters comprising ECM, cell adhesion, and translation components, all of which are implicated in axonal maintenance.

Taken together, these findings show that BDNF drives time-dependent and functionally distinct expression patterns in human motor neurons.

### BDNF increases global rates of transcription in human motor neurons

Next, we explored the transcription kinetics driven by BDNF in human motor neurons. We first measured the effect of BDNF on the direct readout of transcription during SLAM-seq (reads with at least two characteristic T>C mismatches, termed converted reads). Converted reads rose with 4sU labelling time in both control and BDNF-treated motor neurons. BDNF-treated samples had more converted reads at each time point, a difference that reached significance at 6 hours, indicative of increased transcriptional rates (**Fig. 2A**). PCA on converted reads revealed distinct clustering of treatment groups based on time and BDNF treatment (**Fig. 2B)**. To assess transcriptional rate changes driven by BDNF, we used a Monte-Carlo simulation to calculate a new-to-total RNA (NTR) ratio for each gene that met a strict NTR cutoff (90% credible interval < 0.2)(*22*) (**Fig. 2C**). We found that BDNF induces a global upregulation of transcription rate, particularly at 2 and 6 hours, which did not correspond to a rise in steady-state RNA abundance (**Fig. 2D**). This increase in global transcription triggered by BDNF is akin to “hypertranscription,” which is the widespread amplification of transcription during specific cellular contexts, such as during neuronal maturation, cellular growth, and regeneration (*23*). Crucially, these processes are regulated by BDNF in the central and peripheral nervous system (*24*).

**Figure 2.**
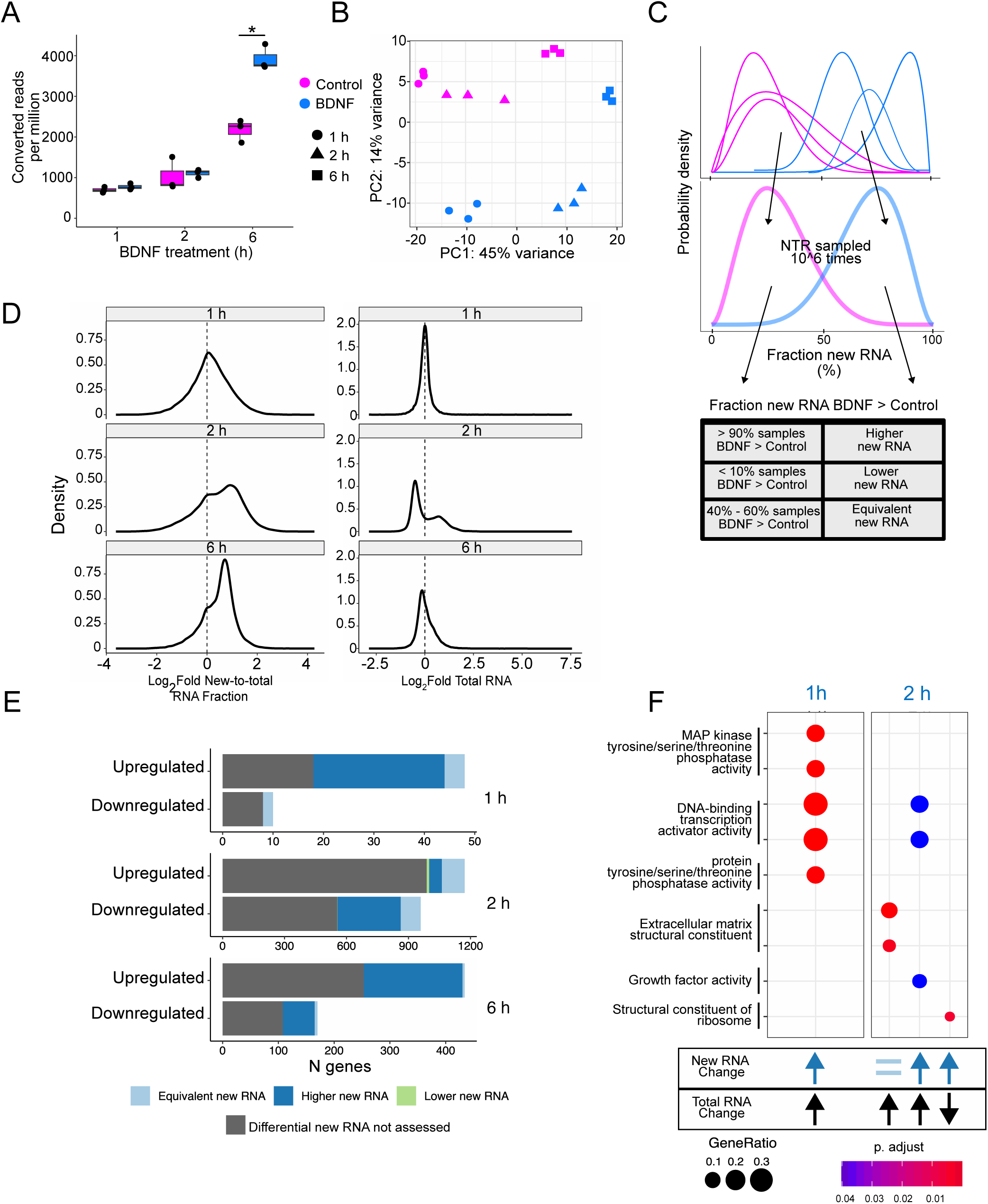
BDNF coordinates global transcription and widespread changes in RNA stability. A) SLAM-seq converted reads with at least 2 characteristic T>C mismatches per million of mapped reads at 1, 2, and 6 h of 4sU treatment in control and BDNF treated i3 LMN. B) Converted reads principal component analysis. C) Schematic of Monte-Carlo sampling approach used to determine condition specific new-to-total RNA ratio (NTR). Beta-distributions on the NTR for a gene for each of N = 3 biological replicates at each time point were estimated by GRAND-SLAM. Condition-specific beta distributions were created by merging the sample beta-distributions with each sample weighted inversely by its variance. Condition NTR was calculated by performing one million samples from each condition’s beta distribution, and differential labelling was determined by comparing the difference between each of the 1 million samples. D) Transcriptome-wide distribution of the log2 fold change of condition specific NTR RNA fraction (left) and the log2 fold change of total RNA. Of genes which met strict NTR cutoff (90% credible interval < 0.2), by 2 h the majority of genes had a higher NTR ratio with BDNF treatment (1 h - 1,392 of 6,169 genes; 2 h - 6,858 of 8,835 genes; 6 h - 8,246 of 8,909 genes). E) Categorization of genes by BDNF effect on total RNA abundance and NTR across treatment times. Stabilised - upregulated total, equivalent or lower new RNA; 1 h - 4 genes; 2 h - 123 genes; 6 h - 4 genes. Destabilisation - downregulated total, higher new RNA; 1 h - 0 genes; 2 h - 306 genes; 6 h - 57 genes, and transcriptionally upregulated - upregulated total, higher new RNA; 1 h - 26 genes; 2 h - 61 genes; 6 h - 177. F) Gene-ontology enrichment for genes categorised in (E). No significant GO terms were found at 6 h treatment time. Significant GO terms using an adjusted p-value < 0.05.

### BDNF signalling alters RNA stability in human motor neurons

We next investigated how BDNF-dependent gene regulation is shaped through transcription and RNA stability mechanisms. We grouped genes by both their change in total RNA levels, as well as a function of newly synthesised RNA to reveal genes with patterns of stabilisation, destabilisation, and transcriptional upregulation (**Fig. 2E; Table S1**). At 1 and 6 hours of BDNF treatment, the majority of upregulated genes showed signs of transcriptional increase with higher new RNA levels (**Fig. 2E; Table S1**). We also identified a group of RNAs that were downregulated, but highly transcribed 2 hours post-BDNF treatment (**Fig. 2E**), which indicates large-scale degradation of RNAs by BDNF after 2 hours of treatment, despite the hypertranscription observed (**Fig. 2D**). Prolonged BDNF treatment in iPSC-derived cortical neurons and mouse GABAergic neurons also lead to widespread downregulation of RNAs (*19*, *25*). These findings suggest sustained treatment of BDNF can drive RNA degradation, although the mechanism and function of this phenomenon are currently unclear.

We then categorised genes by the effect of BDNF on the ratio of total to new RNA across treatment timepoints and performed GO analyses (**Fig. 2F; Table S2**). At 1 hour of treatment, RNA regulation was primarily driven by transcriptional upregulation, affecting MAPK pathway inhibitors, such as *DUSP1*, *DUSP5*, and *DUSP6*, along with early response genes like *ARC*, *FOS*, and the regeneration-associated transcription factor *ATF3* (**Fig. 2F; Table S2**). At 2 hours, transcriptional upregulation also increased the expression of genes associated with growth factor activity, such as the regulators of neurite growth *VEGFA*, *IL11,* and *JAG1* (**Fig. 2F; Table S2**). We also observed a group of genes that showed increases in total RNA abundance, without increases in NTR, suggesting possible RNA stabilisation mechanisms. Notable examples of these genes are those encoding ECM structural proteins (**Fig. 2F; Table S2**). Amidst the robust increase in the newly transcribed RNA ratio for ribosomal genes, these genes are paradoxically downregulated at total RNA level (**Fig. 2F**; **Fig. 1D; Table S2**). In contrast, BDNF has been found to upregulate ribosomal proteins through mTOR-dependent translation in neurons (*26*, *27*). This increased translation of ribosome components, along with evidence that elevated ribosomal flux can destabilise actively translating mRNAs (*28*), may partly explain the concomitant hypertranscription and instability of ribosomal genes.

Thus, we show in human motor neurons that BDNF triggers hypertranscription, while also modulating the expression levels of many genes post-transcriptionally, potentially through changes in RNA stability.

### BDNF drives the expression of regeneration-associated genes and injury-inducible transcription factors

Injury-inducible genes are critical for axonal regeneration (*2*, *29*). Given the role of BDNF in peripheral axon regeneration, we examined its effects on transcriptional programs specifically linked to neuronal injury. We used three high quality compendiums of regeneration-responsive genes (*30–32*) (**Fig. 3A**). We then performed Gene Set Enrichment Analysis (GSEA) on BDNF-induced changes in abundance and newly transcribed RNA in motor neurons (**Fig. 3A,B)**. Strikingly, we found that BDNF-regulated transcripts overlap with regeneration-associated genes (RAGs). At 1 hour, all three datasets showed increased transcriptional rates and RNA abundance (**Fig. 3B,C**), suggesting BDNF rapidly upregulates the transcription of genes critical for neuronal injury response. However, at 2 hours, two gene sets showed a paradoxical pattern similar to what we observed for ribosomal protein genes: total RNA levels decreased while new RNA increased, suggesting active mRNA degradation despite ongoing transcription (**Fig. 2C)**. By 6 hours, total RNA levels were enriched across all datasets, but new RNA enrichment was absent. Altogether, BDNF signalling rapidly upregulates a cluster of genes crucial for neuronal injury response even in the absence of axonal damage. BDNF may therefore increase the responsiveness of motor neurons to axonal injury.

**Figure 3.**
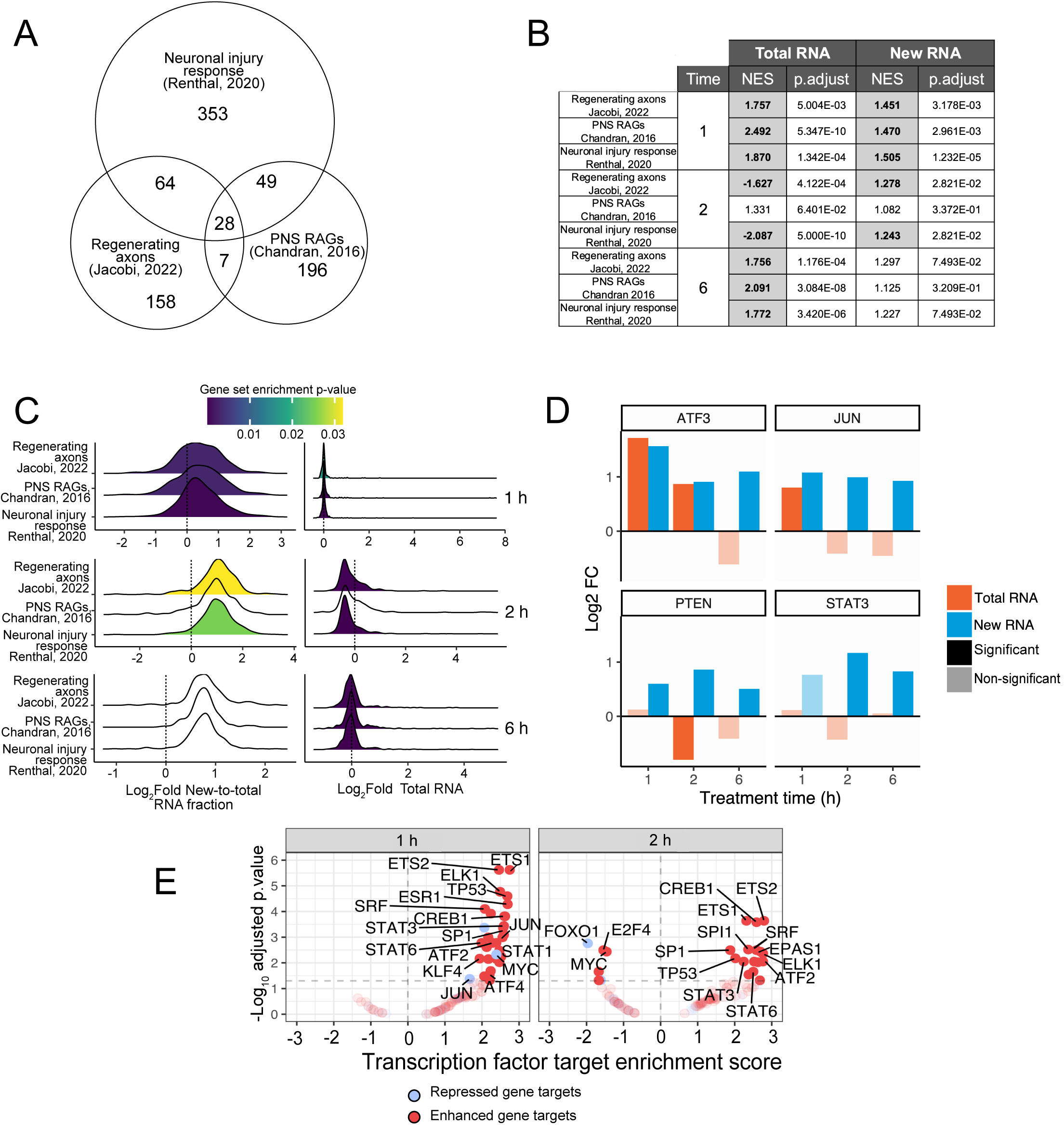
BDNF drives regeneration-associated transcriptional programs in human motor neurons. A) Number of genes and overlap between three different gene sets defining neuronal injury response. Gene sets are originally derived from *Rattus norvegicus* (PNS RAGs, Chandran 2016) and Mus musculus (Regenerating axons, Jacobi 2022 and Neuronal injury response, Renthal 2020) and were first mapped to their human orthologs using gprofiler2 (*66*). B) Normalised enrichment score (NES) and adjusted p-value for the log_2_ fold change of total RNA and NTR on neuronal injury response gene sets in C. Bolded and highlighted enrichments are significant at adjusted p-value < 0.05. C) Distribution of the log_2_ fold NTR and total RNA for axonal regeneration sets. Area under curve coloured: significant; blank: non-significant. Colour refers to the adjusted p-value of the gene-set enrichment analysis of that gene set. NES as listed in B. Significant gene set enrichment analysis (GSEA) with an adjusted p-value < 0.05. D) Total (orange) and new RNA (blue) log_2_ fold change across time for select known regeneration associated genes showing differing effects of BDNF on both abundance and RNA turnover across treatment time. Significant total RNA called with DESeq2 (*62*) and significant new RNA called using Monte-Carlo simulation described in Fig. 2A. E) GSEA transcription factor (TF) regulons from DoRothEA. Red - TF activates target genes; blue - TF represses target genes. Positive TF enrichment scores - target genes upregulated. 6 h - only ETS1 activated targets were significantly enriched (NES = 2.037, adjusted p-value = 0.02069). Significant GSEA with an adjusted p-value < 0.05.

We next investigated the expression patterns of three exemplar RAGs (*ATF3*, *JUN*, and *STAT3)*. We found that these RAGs are highly transcribed by BDNF as shown by a significant increase in NTR across all timepoints (**Fig. 3D**). We found *ATF3* and *JUN* are autoregulated at total RNA levels, since both are only differentially upregulated at 1-2 hours. In contrast, the total levels *of STAT3* did not change amidst the high transcriptional rates (**Fig. 3D**). *PTEN*, a negative regulator of axonal outgrowth, shows increased levels of new RNA, even though we observed a downregulation of its total RNA levels at 2 hours, suggesting the destabilisation of *PTEN* (**Fig. 3D**). These data indicate that BDNF regulates the expression of certain RAGs through RNA stability.

The cell-intrinsic capacity of neurons to respond to axonal insults requires the activation of injury-inducible transcription factors (*2*). To explore which transcription factors are putatively activated by BDNF in motor neurons, we performed GSEA with the DoRothEA database, which curates benchmarked transcription factor-target gene interactions (**Fig. 3E**; Garcia-Alonso et al., 2019). BDNF signalling activated multiple transcription factors at 1 and 2 hours (**Fig. 3E**). We did not detect any enriched TFs at 6 hours except for ETS1. As summarised in **Table 1**, the majority of transcription factors that we identified as BDNF-responsive in motor neurons play a role in neuron-intrinsic regeneration pathways following axonal injury (*2*, *30*). This analysis shows that BDNF signalling potentially fosters the activation of multiple injury-inducible genes sets, which are co-regulated by transcription factors essential for axonal outgrowth and regeneration (*2*, *29–31*).

**Table 1.**
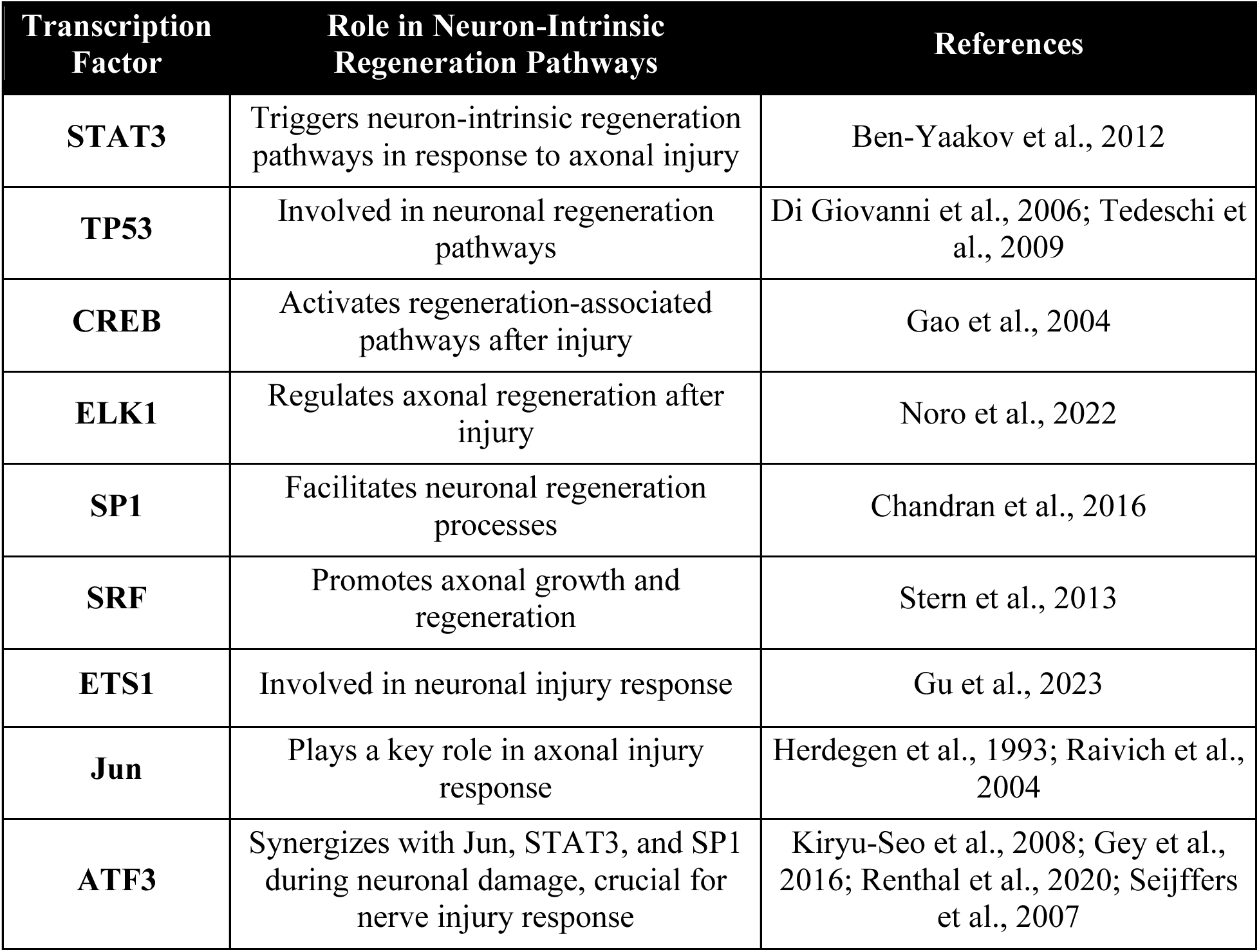
Known transcription factors associated with axonal regeneration identified by presented RNA-sequencing.

Overall, through SLAM-seq and computational analyses of BDNF-responsive genes, we found that BDNF signalling triggers global hypertranscription and controls RNA stability. Furthermore, BDNF rapidly and transiently increases transcription of RAGs and a subset of cell-intrinsic neural repair TFs.

### BDNF reshapes the phosphorylation landscape of cytoskeletal-interacting proteins in human motor neurons

All aspects of axonal repair rely upon precise changes to cytoskeletal dynamics that support the reconstruction of damaged axons. Cytoskeletal restructuring is determined by structural microtubule-associated proteins (MAPs) and actin-binding components (*13*, *34*). BDNF has been shown to phosphorylate components of the cytoskeleton, however its substrates relevant to axonal regeneration are not fully characterized. To address this, we performed tandem mass tag (TMT)-based quantitative phosphoproteomics in i^3^ LMNs upon BDNF incubation (**Fig. 4A; Table S3**). After 1 and 6 hours of BDNF treatment, we confirmed that NTRK2/TRKB is the most phosphorylated protein identified, along with the well-established signalling nodes expected to be activated by BDNF, such as TSC2 and MAP2K1/MEK1 (**Fig. 4B-C**).

**Figure 4.**
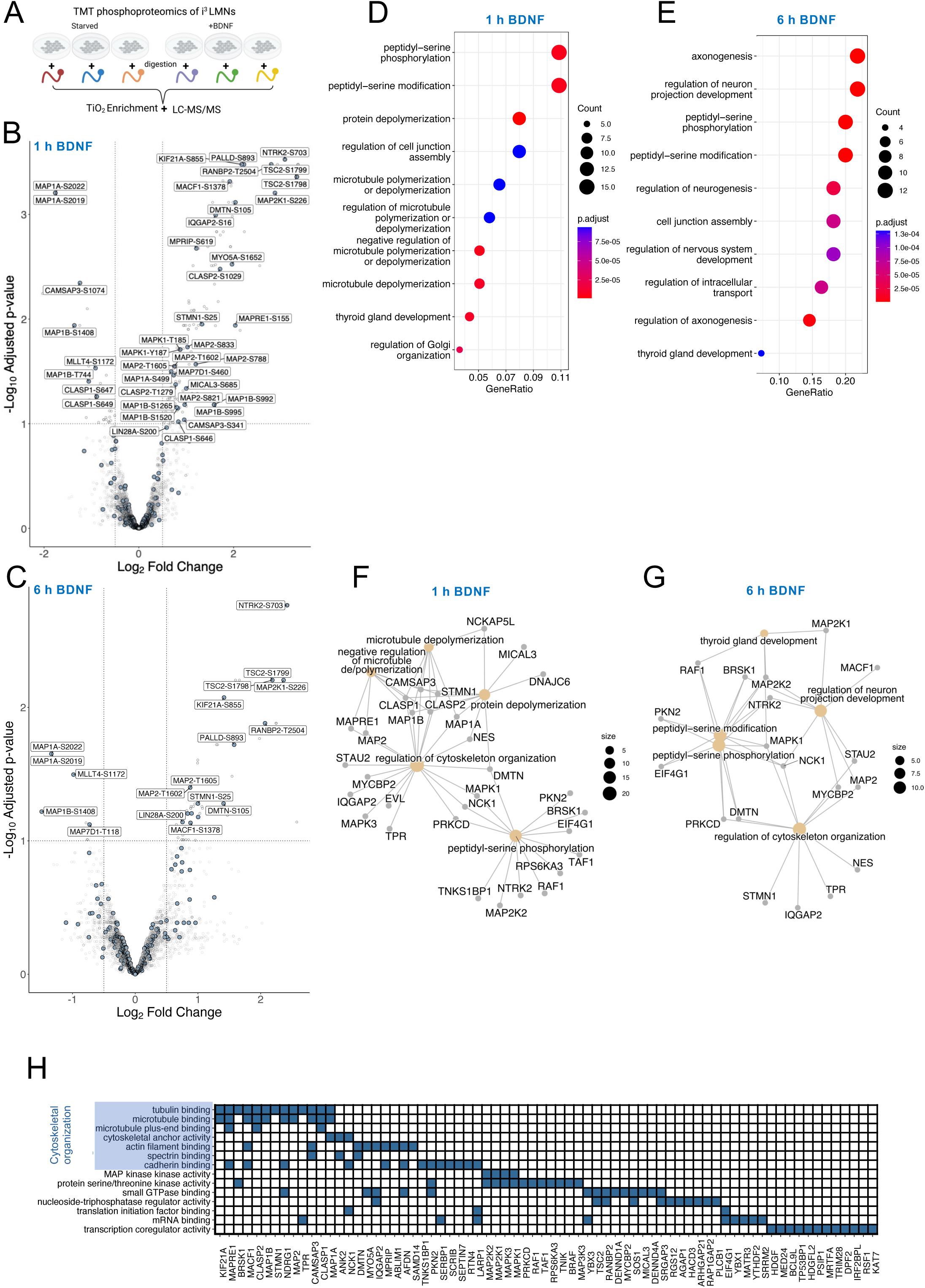
BDNF governs the phosphorylation state of cytoskeletal-binding proteins. A) Schematic of tandem mass tagging (TMT) phosphoproteomics. Unique isobaric tags were added to control, 1 h, and 6 h BDNF-treated DIV7 i3 LMNs. N=3 per condition (No BDNF, 1 h and 6 h 25 ng/mL BDNF). B-C) Volcano plot showing the fold change and adjusted p-values of the phosphorylation sites at 1 h (B) and 6 h (C) after BDNF (25 ng/mL) treatment. Each point represents a phosphosite. Blue dots represent microtubule associated proteins (MAPs) and known downstream targets of BDNF. Significantly differentially phosphorylated sites are labelled with protein name and position in the sequence. Dotted lines represent adjusted p-value 0.1 and log2 fold change +/- 0.5. 148 significant phosphorylation hits at 1 h and 49 hits at 6 h of BDNF treatment (Table S3). D-E) Dotplot showing gene-ontology (GO) enrichment on proteins with significant changes in phosphorylation at 1 h (D) and 6 h (E) after BDNF treatment. F-G) Gene-concept networks depicting the gene-GO relationships for proteins with upregulated phosphorylation at 1 h (F) and 6 h (G) after BDNF treatment. H) Gene-concept heatmap showing the overlap between significantly phosphorylated proteins after 1 h BDNF treatment. The shared gene-ontology terms, which are highlighted in the light blue box, are the GO terms related to cytoskeletal organisation.

Crucially, we discovered striking changes in phosphorylation of several microtubule- and actin-binding proteins downstream of BDNF signalling (**Fig. 4B-C**). The robust enrichment of these proteins in our phosphoproteomic data (**Fig. 4B-G**) is crucial, since their phosphorylation status dictates their function (*15*). Delving deeper into the phospho-regulation of these proteins by BDNF, we found multiple phosphorylation and dephosphorylation events within a single MAP (**Fig. S4A**). For instance, both MAP1A and MAP1B show phosphorylation and dephosphorylation at multiple sites (**Fig. 4B,C; Fig. S4A**). Several of these phospho-events are also time-dependent indicating complex phosphorylation patterns on structural MAPs in response to BDNF (**Fig. 4B,C**).

Other cytoskeletal-binding proteins that are significantly phosphorylated downstream of BDNF include: MAP2, MAP7D, MAPRE1/EB1, MACF1, CAMSAP3, CLASPs, PALLD, DNMT, MICAL3, MLLT4, NCKAP5L, and IQGAP2 (**Fig. 4B-C,F-G; S4A-B**). The functional diversity of these proteins include microtubule-stabilizing and -destabilizing, actin-microtubule crosslinking, microtubule plus and minus end-binding, all of which regulate the cytoskeletal organization of neurons and may be consequential for axonal regeneration (*13–15*). We find that many of these phospho-events are detected proximally or within the actin- and microtubule-binding regions of these proteins, hinting at a regulatory influence of BDNF on the cytoskeletal-binding properties of these proteins (**Fig. S4A-B**). Thus, BDNF promotes the global phosphorylation of cytoskeletal-binding components.

Since we identified multiple cytoskeletal-binding proteins as phospho-substrates of BDNF, we sought to further characterise these events. To this end, we generated a sequence logo (*35*) of the significant phosphopeptides identified, and found that BDNF treatment primarily resulted in phosphorylation of serines followed by a proline at the +1 position (pS-P). Although not as pronounced as pS-P, the basophilic RXRXXpS/T (X = any amino acid) phospho-motif canonical to the AGC superfamily kinases (*36*) is also evident in our motif analysis (**Fig. S4C**). Proline-directed kinase families include ERK, mTOR, and cyclin-dependent kinases, which are known downstream effectors to TrkB activation. When we focused our analysis on proteins with significant proline-directed serine phosphosites, we found an enrichment of cytoskeletal-binding proteins (**Fig. S4D**). We validated by western blotting two proline-directed BDNF-dependent phosphorylation events on STMN1 (S25), a microtubule destabilising enzyme, and Lin28a (S200), an RNA-binding protein (RBP) known to enhance axonal regeneration (*37*)(**Fig. S4E-H**). Both phosphorylation sites are driven by BDNF and inhibited by the addition of U0126, a well-established ERK inhibitor, indicating that BDNF triggers the phosphorylation of at least these two sites through ERK in human motor neurons (**Fig. S4E-H**).

### BDNF phospho-substrates are enriched for regulators of cytoskeletal dynamics and axonogenesis

We next performed unbiased functional enrichment analyses on the significant BDNF-dependent phospho-events in i^3^ LMNs. At both 1 and 6 hours of BDNF treatment, we reveal an overrepresentation of the microtubule polymerization and depolymerization pathways (**Fig. 4D-E**). Additionally, pathways related to axonogenesis, neuron projection, and regulation of intracellular transport processes, all of which depend on cytoskeletal organization, are highly represented among BDNF substrates (**Fig. 4D-E**). Next, we created gene-concept networks of upregulated phospho-events to better illustrate the enriched functional networks associated with BDNF phospho-signalling (**Fig. 4F-G**). The gene-concept networks revealed several signalling hubs with well-characterised functions on the neuronal cytoskeleton (**Fig. 4F-G**). These hits include BRSK1, which regulates neuronal polarity through Tau phosphorylation (*38*), and the kinase RPS6KA3/RSK2, which has been reported to enhance axonal regeneration (*39*). Additional hits were MYCBP2, an atypical E3 ligase important for regulation of Wallerian degeneration (*40*), and NCK1, which was found to stabilise neuronal actin dynamics (*41*). We also identified an understudied microtubule plus end-binding protein, NCKAP5L (*42*), as a novel phospho-substrate of BDNF (**Fig. 4F-G**). Thus, our unbiased computational approaches reveal that BDNF regulates not only the phosphorylation of many cytoskeletal components, but also of molecular pathways implicated in cytoskeletal dynamics.

Other pathways enriched by BDNF treatment are the regulation of MAPK/ERK signalling, translation initiation factor binding, mRNA binding, and transcription coregulation activity (**Fig. 4H**). The ALS-related RBP MATRIN3, known to regulate RNA stability, is also highly phosphorylated upon BDNF treatment (**Table S3)**. Other RBPs that control mRNA stability, such as STAU2, YBX1 and YTHDF2, were also identified as phospho-substrates of BDNF. Of note, YTHDF proteins are also known positive regulators of axonal regeneration via regulating RNA stability (*43*). Moreover, the ERK-dependent phosphorylation of YTHDF2 at serine 39, detected in our dataset (**Table S3)**, facilitates its RNA degradative function (*44*). These RBPs impact RNA metabolism in neurons and are potentially involved in regulating the post-transcriptional tuning of RNA levels induced by BDNF described herein; although more work is required to validate this hypothesis (**Fig. 2**.)

Taken together, these data indicate that the phosphorylation targets downstream of BDNF in human motor neurons are highly enriched for cytoskeletal components, as well as various signalling pathways involved in controlling neuronal morphology and axonal regeneration.

### BDNF phosphorylation events are enriched for RSK and S6K kinase motifs

Following up on our TMT phosphoproteomics experiment, we used a kinase prediction tool based on large-scale combinatorial synthetic peptide libraries, which can identify the substrate sequence specificity of 300+ kinases (*45*). Using the significant phosphorylation events driven by BDNF as inputs, we found that AGC-family kinases, RSK/p90RSK and S6K/p70S6K, are predicted to be highly enriched kinases downstream to BDNF activation (**Fig. 5A,B**).

**Figure 5.**
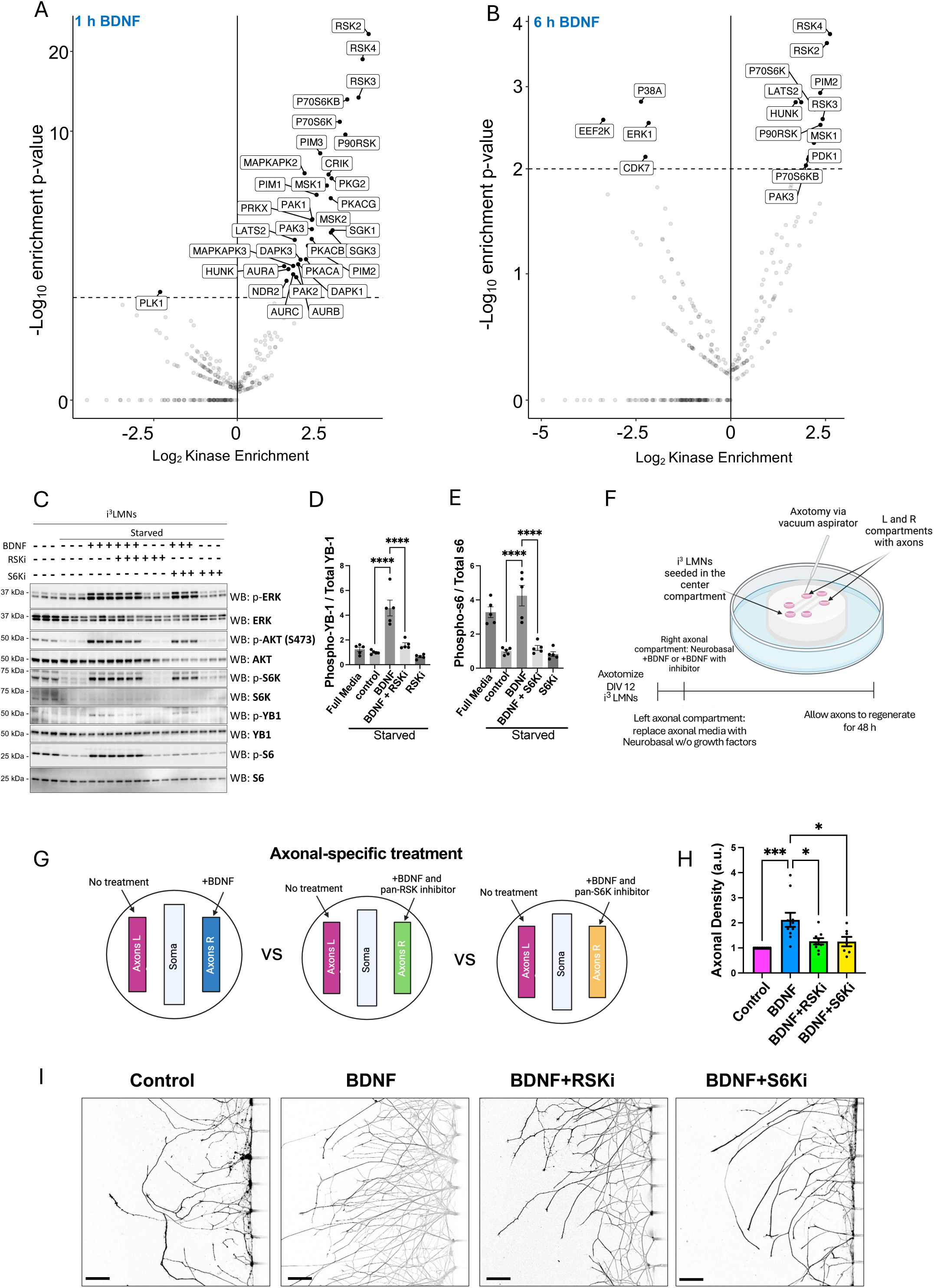
S6K and RSK activation is required for BDNF-dependent axonal regeneration. A-B) Kinase enrichment at 1 h (A) and 6 h (B) after BDNF treatment based on the sequence of differentially phosphorylated sites. Dotted line shows significant enrichment at p-value < 0.01. C) Western analysis of RSK and S6K kinase signalling after BDNF treatment (1 h; 25 ng/mL), or with co-treatment of S6Ki (PF-4708671; 10 μM) or RSKi (LJI308; 1 μM). D-E) Quantification of the western blot analysis shown in (C). N=5. F) Schematic of microfluidic chambers and aspiration-based axotomy. G) Schematic of the treatment conditions of axonal regeneration assay. H) Quantification of axonal density after axotomy and 48 h recovery with or without BDNF (50 ng/mL) or co-treatment with RSK inhibitor LJ308 (1 μM) or the S6K inhibitor PF-4708671 (10 μM). N = control, 10; BDNF 10; BDNF+RSKi, 9; BDNF + S6Ki, 7. I) Representative confocal images of axonal density of i^3^ LMNs cultured in tripartite MFCs 48 h post-injury. LMNs were immunostained with β3-tubulin. Western blot analyses and axonal regeneration experiments were performed using i^3^ LMNs derived from at least two independent differentiations. Replicates consist of one MFC. One-way ANOVA. Error bars: SEM. p value: ∗ = < 0.05; ∗∗ < 0.01; ∗∗∗ < 0.001; ∗∗∗∗ < 0.00001.

RSKs and S6Ks respond to growth factors via the MAPK and PI3K/AKT pathways (*36*). Indeed, BDNF robustly activates RSKs and S6Ks kinases as assessed by the phosphorylation of two well-known targets of these pathways, YB-1 and S6, respectively (**Fig. 5C-E**). Furthermore, YB-1 and S6 phosphorylation are selectively inhibited by a pan-RSK and a pan-S6K inhibitor, respectively. This finding indicates these inhibitors do not cause cross-inhibition (**Fig. 5C-E).** Inhibition of ERK after BDNF treatment led to decreased activation of both RSKs and S6Ks (**Fig. S5A-C**), whereas AKT activity is only required for the S6K signalling arm, but not for RSK, in human motor neurons (**Fig. S5D-F**), indicating that these kinases are activated by BDNF via a canonical pathway in human motor neurons.

### Axon-specific RSK and S6K activity is required for BDNF-dependent axonal regeneration

We next investigated whether RSK and S6K kinase activation is critical for BDNF function in axonal regeneration in human motor neurons. To test this, we cultured human motor neurons in tripartite microfluidic chambers (MFCs) and performed axonal regeneration assays with simultaneous treatment with BDNF, along with pan-RSK or pan-S6K inhibitors (**Fig. 5F-G**). Because the two axonal compartments of these MFCs are fluidically isolated from each other and from the somatic compartment, we were able to ascertain the impact of axon-specific treatment of BDNF in one axonal compartment and the simultaneous inhibition of RSK or S6K in the other, in the same motor neuron culture (**Fig. 5F-G**). We axotomized human motor neurons with strong aspiration and allowed the axons to regenerate for 48 hours. As shown in **Fig. 5H-I**, axon-specific treatment with BDNF increased axonal regeneration. This upregulation by BDNF is blocked when either RSKs or S6Ks are blocked only in axons (**Fig. 5H-I**). Thus, our data suggest that the upregulation axonal repair by BDNF depends on the axon-specific activation of RSKs and S6Ks. Furthermore, these experiments functionally link the BDNF-dependent phosphorylation of the substrates reported in Fig. 4 and Fig. S4 to axonal regeneration through RSKs and S6Ks.

### Axonal BDNF enhances axonal regeneration of human motor neurons through localised ERK activation

Lentiviral transduction of TrkB *in vivo* enhances the axonal regeneration of corticospinal neurons though ERK signalling (*6*). Moreover, RSK and S6K activity are dependent on ERK upon BDNF treatment. Thus we next explored the requirement for axon-specific activation of ERK by BDNF on axonal regrowth post-injury. To this end, we performed axotomy experiments using i^3^ LMNs cultured in tripartite MFCs (**Fig. 6A**). We found the axon-specific effect of BDNF in fostering regeneration is blocked upon ERK inhibition (**Fig. 6B-C; Fig. S6A-B**). These results are in line with our phosphoproteomic analysis showing an enrichment of proline-directed motifs, potentially directed by ERK, on cytoskeletal-binding proteins upon BDNF treatment (**Fig. S4A-C; Fig. S4G-H)**. In contrast to ERK inhibition, the axonal-specific inactivation of AKT by Ipatasertib does not affect BDNF-dependent regeneration (**Fig. 6B,C; Fig. S6C-D**). Deletion of PTEN, which negatively regulates the PI3K/AKT pathway, is well-established to foster axonal regeneration (*46*). We therefore asked whether AKT signalling, when inhibited specifically in the soma, would affect the injury response of motor neurons. Indeed, inhibition of AKT, as well as ERK, in the somatic compartment hindered axonal regeneration post-injury (**Fig. 6D-F**), demonstrating that somatic activation of both ERK and AKT are required for nerve regeneration. Both ERK and AKT activity are rapidly upregulated by BDNF, however, the activity of ERK is sustained longer than that of AKT (**Fig S6E-G**). Furthermore, we observed quantitative differences in the potency of activation of ERK and AKT via BDNF in the soma and axonal compartments (**Fig. S6H-J**). Although both kinases are activated in the soma and axons, the activation of ERK is more pronounced in the axon, whereas the AKT is more active in the soma (**Fig. S6H-J**). This result potentially contributes to the compartment-specific effects of inhibiting ERK and AKT with respect to BDNF-dependent axon regeneration.

**Figure 6.**
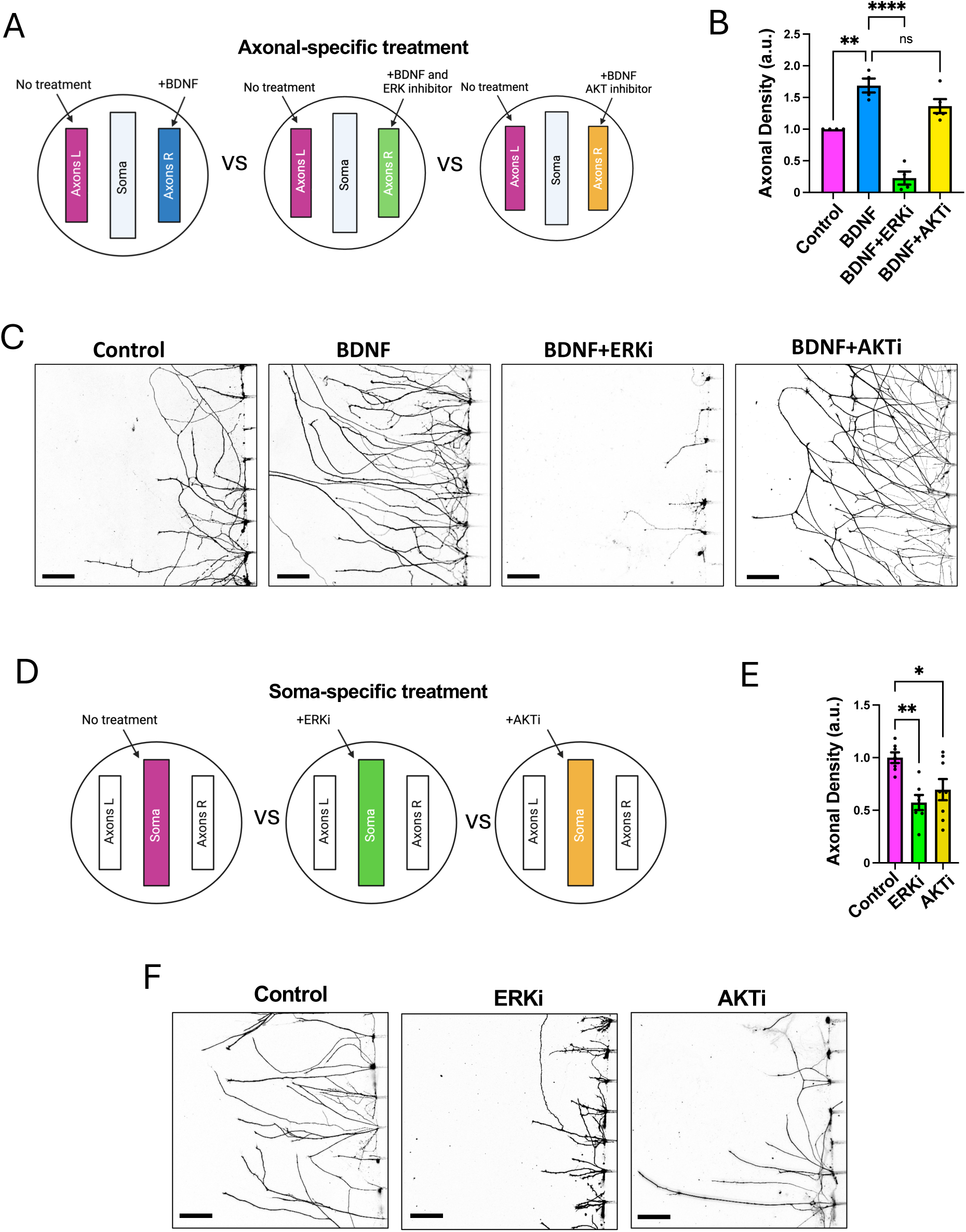
Axon-specific activation of ERK by BDNF enhances regeneration. A) Schematic of axotomy experiments testing the axon-specific treatment of the ERK inhibitor U0126 and the AKT inhibitor ipatasertib on BDNF enhanced regeneration. B) Axonal density after axotomy and 48 h recovery with or without BDNF (50 ng/mL) or co-treatment with U0126 (20 μM) or ipatasertib (0.1 μM) only in axons. N = BDNF, 4; BDNF+ERKi, 4; BDNF+AKTi, 5. C) Representative confocal images of axons from B. i^3^ LMNs were immunostained with β3-tubulin. Scale bars: 50 μm. D) Schematic of axotomy experiments testing the soma-specific treatment of ERK inhibitor U0126 and AKT inhibitor ipatasertib on BDNF enhanced regeneration. E) Axonal density after axotomy and 48 h recovery with or without U0126 (20 μM) or ipatasertib (0.1 μM) treatment of the somatic compartment. N= ERKi, 7; AKTi, 8. All axonal regeneration experiments are performed with neurons derived from at least two independent differentiations. Replicates consist of one MCF. One-way ANOVA. Error bars: SEM. p value: ∗ = < 0.05; ∗∗ < 0.01; ∗∗∗ < 0.001; ∗∗∗∗ < 0.00001.

Our findings demonstrate that BDNF signalling promotes axonal regeneration by locally activating ERK-S6K-RSK pathway specifically within axons, independent of PI3K/AKT. Thus, BDNF-driven axonal repair relies on a specific subset of kinases, as well as their activation within distinct neuronal compartments.

## Discussion

A central question in the field of axonal regeneration is how neurotrophins enhance nerve repair. Efficient nerve regeneration depends on a complex interplay between intrinsic programs and extrinsic factors. In this study, we reveal that BDNF promotes axonal repair and acts as a signalling hub for multiple neuron-intrinsic regeneration programs in human motor neurons.

Optimal RNA expression relies on the timely coordination of transcription, decay, and stability (*47*). Transcriptome tuning is particularly crucial for injury response and axonal repair (*2*, *48*). Through SLAM-seq, our findings demonstrate that BDNF exerts highly complex effects on RNA stability in human motor neurons. Unexpectedly, we also found that BDNF drives hypertranscription, which is not reflected in steady-state RNA levels. This suggests that BDNF calibrates levels of specific RNAs through stabilisation and/or degradation, amidst increased global transcriptional rates. Several RBPs known to regulate RNA stability are top phospho-substrates of BDNF in human motor neurons, which may underlie this phenomenon. Moreover, BDNF regulates the DICER complex and the biogenesis of myriad microRNAs (*49*). Thus, SLAM-seq revealed a novel regulation of RNA metabolism by BDNF previously unobserved using traditional bulk RNA-seq strategies.

An important question arising from our work is the functional significance of hypertranscription and its relevance to axonal regeneration. Hypertranscription has been shown to facilitate tissue regeneration outside of the nervous system and to accommodate for biosynthetically demanding cellular processes (*23*). In response to axonal injury, adult neurons revert to embryonic transcriptional states (*50*) and hypertranscription is more prevalent in developing neurons (*23*). Thus, performing SLAM-seq after axonal injury has the potential to address whether hypertranscription is a widespread injury-induced response. Additionally, if other neurotrophic factors regulate transcriptional responses through hypertranscription warrants further study. Indeed, there is a paucity of data on the impact of neurotrophic factors on neuronal mRNA stability and decay, which needs to be addressed to fully understand the impact of these neuronal growth factors on neuronal transcriptomes. This is crucial given that stabilisation of RNAs is an emerging paradigm controlling the subcellular localization of RNAs in neurons (*8*).

The cell-intrinsic capacity of neurons to sustain axonal outgrowth and regeneration require the activation of specific TFs (*2*, *51*). Amidst the profound changes in the transcriptome driven by BDNF, we found that many of the TF-target gene interactions governed by BDNF are enriched for TFs linked to axonal regeneration. We speculate that these TFs function synergistically to orchestrate genes upon BDNF activation (*52*), fostering nerve repair. Moreover, we find that BDNF rapidly initiates the transcription of injury-inducible RAGs in human motor neurons. Thus, BDNF unlocks the basal cell-intrinsic transcriptional capacity of motor neurons, priming them to respond more aptly to axonal insults. In line with this, genetic manipulations known to improve axonal repair, such as *PTEN* deletion, induce the expression of RAGs independent of axonal injury (*31*). Future transcriptomic work investigating the effect of BDNF after axonal denervation is warranted to ascertain whether BDNF is also able to transcriptionally downregulate neuronal “death” signals post-injury.

Another aspect that dictates the efficacy of neuron-intrinsic injury response is the modulation of cytoskeletal dynamics (*13*, *14*). We demonstrate that BDNF reshapes the phosphorylation landscape of cytoskeletal-interacting proteins, particularly of structural MAPs and actin-binding proteins. Many of the identified phosphosites are positioned within or adjacent to the cytoskeleton-binding domain, suggesting that BDNF regulates the interaction of these proteins with the cytoskeleton and their function through phosphorylation (*15*). The global phosphorylation of cytoskeletal-interacting proteins by BDNF may thus underlie the capacity of BDNF to modulate the architecture of multiple neuronal compartments and to foster axonal regeneration. The functional relevance of these novel phosphorylation events to axonal repair and cytoskeletal dynamics require further exploration, and novel experimental approaches capitalising on reconstituted MAPs and microtubule systems could be an appealing step forward (*34*).

Our kinase prediction analysis revealed RSKs and S6Ks substrates are enriched after BDNF treatment. We further show that axon-specific activation of ERK, and its downstream kinases RSKs and S6Ks are crucial for the axonal regeneration driven by BDNF in human motor neurons. Congruent with our observations, RSKs were previously found to be critical for axonal regeneration (*39*, *53*). Interestingly, modulating multiple kinases is a strategy at the core of combinatorial therapies to enhance axonal regeneration. Nevertheless, the interrelationship between injury-responsive kinases with respect to boosting regeneration is, however, complex and may yield paradoxical effects (*48*). It is therefore not completely surprising that S6K1-specific inhibition was shown to increase axon outgrowth (*54*), in contrast to pan-inhibition of S6Ks, which hinder axonal outgrowth, as we report here. Our data also show that AKT activates RSKs in motor neurons, yet axon-specific activation of AKT by BDNF can be bypassed in terms of its effects in axon repair. Overall, we found that the compartment-specific activation of a subset of kinases by BDNF confers distinct outcomes on axonal repair in human motor neurons.

ERK is activated after peripheral nerve injury to promote axonal regeneration in mice (*6*, *55*) and has well-established roles in cytoskeletal reorganization in axons. RSKs and S6Ks also have a direct impact on microtubule and actin dynamics by phosphorylating specific cytoskeletal components, as inferred from a limited number of studies in mitotic cells (*56*, *57*). Whether the activation ERK-RSK-S6K pathway by BDNF is responsible for the changes in the phosphorylation landscape of cytoskeletal-binding proteins and the modulation of the neuronal cytoskeleton is worth exploring in future studies.

Neuron-intrinsic pathways boost axonal regeneration through the coordination of transcriptional programs, precise kinase activation patterns, and changes in cytoskeletal dynamics (*3*, *13*, *48*). Here, we uncover that BDNF, an extracellular ligand, orchestrates various neuron-intrinsic programs critical for axonal repair. Our data may have substantial implications for the fine-tuning of neurotrophin-based therapies, and novel combinatorial pharmacological interventions targeting both intrinsic and extrinsic regeneration pathways (*58*), geared towards enhancing axonal viability after traumatic injuries and neuropathies.

## Materials and Methods

### Human iPSC cell culture

A human induced pluripotent stem cell (iPSC) line (WTC11), stably expressing a doxycycline-inducible hNIL construct and CRISPR/Cas9 was used in this study. Briefly, iPSCs were seeded on tissue culture dishes coated with 120-180 μg/ml Geltrex (Thermo Fisher). Cells were maintained in iPSC medium containing Essential 8 Flex Medium (E8F, Thermo Fisher), E8 Flex supplement (Thermo Fisher), and 100 U/ml Pen/Strep and maintained in a 37 °C, 5% CO2 incubator. Medium change was performed every 1-2 days. Passage of cells were performed with Stem Pro Accutase (Thermo Fisher) for 1-2 min at 37 °C. Accutase was removed by centrifugation at 300 g for 5 min, and cells were reseeded in iPSC media supplemented with 10 μM ROCK inhibitor (Y-27632, Selleckchem) to facilitate survival. ROCK inhibitor was removed after 24 h.

### i^3^ LMN differentiation from human iPS cells

Differentiation of human iPSCs was performed as previously reported. Briefly, iPSC induction was achieved using induction medium containing DMEM/F12, GlutaMAX supplement medium (Thermo Fisher), MEM non-essential amino acids (Thermo Fisher), N2 supplement (Thermo Fisher), 0.2 mM compound E, 2 μg/ml doxycycline, Pen/Strep and 10 μM Y-27632. After induction for 48 h, cells were dissociated from plates by Accutase and reseeded on poly-D-lysine (PDL)/laminin-coated tissue culture dishes or microfluidic devices in induction medium supplemented with 1 μg/ml laminin. After 24 h, induction medium was replaced by motor neuron medium containing Neurobasal (Thermo Fisher), B27 Plus supplement (Thermo Fisher), N2 supplement, Culture One supplement (Thermo Fisher), 1 μg/ml laminin, 2 μg/ml doxycycline, and Pen/Strep. Medium change was performed every 1-3 days.

### Western blot analyses

Samples for western blot were denatured in lithium dodecyl sulfate (LDS, pH 8.4, NuPAGE) containing 100 mM dithiothreitol (DTT, Sigma) by boiling at 99 °C for 10 min. The denatured samples were separated on 4-12% Bis-Tris Midi gels (NuPAGE) according to the manufacturer’s instructions. Proteins were transferred to polyvinyl difluoride (PVDF) membranes and immunoblotted using antibodies (1:1,000 or 1:500 in blocking buffer; 1X PBS in 3% milk or bovine serum albumin (BSA)) as specified in the reagent table. For microfluidic chamber western blots, axonal compartments from 3-4 chambers are combined for axonal-enriched lysates.

### Immunofluorescence

Cells were fixed with 4% paraformaldehyde (PFA) in phosphate-buffered saline (PBS) at room temperature for 10 min. Fixed cells were washed with PBS three times. For permeabilization and blocking, cells were incubated with 0.1% saponin, 3% BSA in PBS at room temperature for 1 h. Permeabilized cells were then incubated with 0.05% saponin, 3% BSA in PBS supplemented with antibodies specified in the reagent table overnight at 4 °C. Cells were then washed with PBS 3 times before incubation with AlexaFluor-conjugated secondary antibodies (Thermo Fisher) at room temperature for 1 h. Cells were then washed with PBS again 5 times before confocal imaging. Images were captured using an inverted Zeiss LSM 980 microscope using a 63x oil-immersed objective.

### Fabrication of tripartite microfluidic devices

The three-compartment microfluidic devices were fabricated using polydimethylsiloxane (PDMS). Moulds with 500 μm long microgrooves were then cured and attached to a PDL-coated Willco glass dishes. After washing with water, the chambers were coated with 50 μg/ml laminin and 0.8% embryonic stem (ES) BSA in DMEM/F12 medium. Prior to seeding, the chambers were washed once with Neurobasal. Differentiated motor neurons (DIV 0 post-differentiation) were seeded into the middle ‘somatic’ compartment with induction media then replaced with motor neuron media after 24 h. Neurons were fed by replacing the motor neuron media every 1-2 days until DIV 12-14 prior to axotomy and imaging experiments.

### Axotomy and axonal regeneration of human motor neuron

Differentiated i^3^ LMNs seeded in three-compartment MFCs were subjected to axotomy at DIV12-14 post-differentiation. Axons were severed by strong aspiration from the axonal compartment and examined under a light microscope to ensure that all axons were severed. MN media or starvation media with drugs indicated in figure legends were then added to the axonal compartment. After 48 h, cells were fixed and processed for immunofluorescence using an anti-b3 tubulin antibody to visualise axons. Confocal microscopy was then performed using a Zeiss LSM 980 (20X objective). For each tripartite microfluidic chamber, at least five images from each of the axonal compartments were acquired. Only fields with axonal regrowth were captured. Using ImageJ, axonal images were binarized and the total area of axons within the region of interest was calculated. Axonal images were then averaged to determine the total area of axonal regrowth in each axonal compartment. For statistical analysis, each microfluidic device was considered as a replicate and neurons from at least 2 differentiation rounds were used to account for batch-to-batch variability of differentiation.

### LC-MS/MS analysis

For analysis on whole cell and phospho-proteomics, cell lysates were digested using the S-trap (ProtiFi) protocol according to the manufacturer’s instructions. Briefly, proteins were reduced using TCEP (10 mM, 30 min, 56 ℃) followed by alkylation using iodoacetamide (20 mM, 30 min, under dark). Samples were acidified prior to transfer to the S-trap micro. Proteins were digested in the trap using a trypsin/Lys-C mix. Peptides were cleaned using Sep-Pak C18 minicolumns using manufacturer’s instructions. Cleaned peptide samples were then enriched for phospho-peptides using Titansphere TiO_2_ 5 µm (GL Sciences) beads. Resultant phosphopeptides were then TMT-labelled and a pool of 5% of each TMT-labelled phosphopeptide samples was created and analysed with MS to balance the input amounts for each sample. A pool of equal amounts of each labelled phosphopeptide sample was created and fractionated using a Vanquish HPLC at high pH into 48 fractions. These fractions were dried down and resuspended in 0.1% formic acid (Thermo Fisher) prior to MS analysis. Fractionated phosphopeptides were analysed on a Orbitrap Fusion Lumos Tribrid mass spectrometer interfaced with a 3000 RSLC nano liquid chromatography system. Roughly 1 mg of each sample were loaded onto a 2 cm trapping column (PepMap C18 100A – 300 mm, Thermo Fisher) at 5 µl/min flow rate using a loading pump and analysed on a 50 cm analytical column (EASY-Spray column, 50 cm x 75 μm ID) at 300 nl/min flow rate, that is interfaced to the mass spectrometer using Easy nLC source and electro sprayed directly into the mass spectrometer. A linear gradient between solvent A consisting of 0.1% formic acid in water and solvent B consisting of 0.1% formic acid in 99.9% acetonitrile (Thermo Fisher) was used. Firstly a gradient of 3% to 30% of solvent B was used at a 300 nl/min flow rate for 105 min and 30% to 40% solvent B for 15 min and 35% to 99% solvent B for 5 min, which was maintained at 90% of B for 10 min and washed the column at 3% solvent B for another 10 min comprising a total of 140 min run with a 120 min gradient in a data dependent acquisition (DDA).

### Mass spectrometry data analysis and bioinformatics analysis

For analysis on whole cell and phospho-proteomics, resultant MS data files were searched using MaxQuant (ver. 2.2.0) against a human database (Uniprot SwissProt database dated February 2022). The search was performed using the default settings with the fixed modification of TMT labels at the MS2 level and the variable modification of Phosphorylation on STY residues. The produced matrix of phosphopeptide abundances was processed using R (ver. 4.1.3) to normalise the data. Missing values were imputed using the Mice R package.

### SLAM-seq sequencing

SLAM-sequencing was performed on i^3^ LMNs following protocols adapted from Herzog et al. (2017)(*17*). Three biological replicates of i^3^ LMNs were treated with 100 µM 4sU (Control) + labelling medium or 100 µM 4sU + BDNF (BDNF condition) labelling medium on day 7 for 1 h, 2 h, and 6 h. Cells were washed with PBS and RNA was extracted using the Qiagen RNeasy isolation and purification kit. RNA concentration was measured, and after equilibrating the amount of RNA in each sample, iodoacetamide (10 mM), Na_2_HPO_4_ (pH 8, 50 mM), and DMSO (50% v/v) were added to each sample and incubated at 50 °C for 15 min to facilitate the thiol-alkylation of 4sU. Samples were processed on RNeasy columns to re-isolate RNA. Sequencing libraries were prepared using the KAPA HyperPrep Kit with RiboErase kit (Roche). Samples were sequenced at 2 x 100 bp on an Illumina NextSeq 2000 machine.

### Total RNA differential gene expression

Sequencing files were trimmed for adapters using Fastp with the parameter “qualified_quality_phred: 10”(*59*) and quality was assessed using fastqc (*60*). Quality control examination of sequencing data led to exclusion of one of the four biological replicates being called an outlier for both control and BDNF treated samples, leading to a final n = 3 biological replicates for both controls and BDNF-treated samples. Samples were aligned to the Genome Reference Consortium Human Build 38 (GRCh38) genome using hierarchical indexing for spliced alignment of transcripts - 3 nucleotides (HISAT-3N) (*61*) and quantified using FeatureCounts using gene models from GENCODE v34. For the differential expression analysis of the total RNA, all samples were run using the standard DESeq2 (*62*) workflow in R. An adjusted p-value < 0.1 and an absolute log_2_ fold cutoff of 0.75 was used to determine up and downregulated genes on the total RNA level. For differential expression of the previously published RNA-seq(*25*) dataset, consisting of 2 hours BDNF treated H9 human embryonic stem cell-derived GABAergic inhibitory neurons, we downloaded the count and metadata matrices provided on ArrayExpress (accession no. E-MTAB-6975) and input into the standard differential expression workflow. For the gene-set enrichment analysis on the various axonal injury gene sets for input, to account for variance, we used ‘apeglm’ to shrink log_2_ fold changes prior to the gene-set enrichment analysis.

### New-to-Total RNA labelling analyses

Globally refined analysis of newly transcribed RNA and decay rates using SLAM-seq (GRAND-SLAM) was used to compute the proportion and corresponding posterior distribution of new and old RNA for each gene(*18*). To test the difference in the fraction of newly transcribed RNA in BDNF and control conditions, we used a Monte-Carlo sampling model. For each gene, we first selected samples where the range of the upper and lower estimate of the newly transcribed ratio was less than 20%. If a gene did not have at least 2 of 3 samples passing this threshold, we did not assess whether the gene was differentially transcribed. Next, we generated a condition probability density for the newly transcribed ratio by generating a beta distribution from all passing samples per condition and inversing weighting each sample by its variance. Then we sampled 1 million times from each condition-specific beta distribution and compared whether the sample was greater in BDNF versus control. Genes were categorised by comparing the fraction of the 1 million samples from each beta distribution where the estimate of new RNA in BDNF condition was higher than the estimate sampled from the control. If 90% of the sampled new RNA estimates was higher in the BDNF compared to Control, then a gene was categorised as ‘bdnf_higher_new_rna’, if less than 10% of samples was higher in BDNF, then a gene was categorised as ‘bdnf_lower_new_rna’, if the BDNF estimate was higher in 40% to 60% of the sampled new RNA estimates, the gene was categorised as ‘bdnf_equals_control’. To control for potential confounding effects of dropout, where highly labelled RNA is underrepresented in metabolic labelling sequencing due to misalignment during analysis (*63*), we used specialized alignment software (HISAT-3N).

### Total gene expression clustering

Genes were hierarchically clustered by the total RNA log_2_ fold change across time points using ‘ward.D2’ method from the R stats package, and the optimal number of clusters was chosen based on a visual assessment of the cluster plot. Specifically, the plot was examined to identify the number of clusters that demonstrated the highest level of cohesion among members while maintaining a distinct separation from the other clusters.

### Transcription Factor Enrichment Analysis

Gene regulatory networks (GRN) from DoRothEA TF database (*33*) were first filtered to include only TF-target interactions with confidence levels A. These are the TF-target interactions with the highest level of confidence with multiple lines of evidence supporting the directional interaction, including ChIP-seq, TF binding motif on promoter, manual curation by experts in reviews, or supported in at least two curated resources. We also filtered for GRN with a minimum set size of 5 genes. DoRothEA reports both positive and negative regulatory targets and these GRN were kept as separate gene sets. To control for gene expression significance, we used as input to the GRN all genes ranked according to their p-log10 transformed pvalue signed by the direction of change. GSEA was performed using the ‘fgsea’(*64*) package in R. Full analysis available in Table S3. Significant GSEA was called with an adjusted p-value < 0.05.

### Gene-ontology enrichment analyses

All gene-ontology enrichment was performed in R using the clusterProfiler (*65*) package. In the GO analysis combining total and NTR information, we did not include the GO genes which were either “not significant” on total RNA or where we could not accurately assess differential NTR based on the criteria laid out in the methods section on NTR labelling. To ensure that enriched GO terms would pick up pathways specific to the effects of BDNF treatment and not terms related to tissue-specific expression, only genes which were detected by RNA-seq in the i^3^ LMNs were used as the background gene set. Full GO term analyses are available in Table S3.

### Gene set enrichment of neuronal regeneration datasets

We identified three gene sets associated with neuronal regeneration. From the Jacobi 2022 paper, we used the 306 genes used to create a ‘composite rag score’ which were selectively expressed in regenerating retinal ganglion found in Table S4 of the paper (*31*). From the Chandran 2016 paper, we selected genes from the ‘magenta’ module which were identified as being upregulated after peripheral nerve injury from Table S2 of the paper (*30*). From the Renthal 2020 paper, we selected the genes which were common injury genes across neuronal subtypes in Table S4 of the paper (*32*). We then performed GSEA against these gene sets using both the change in transcription rate, as measured by the log_2_ fold change of the NTR between BDNF and control, and the change in RNA abundance, as measured by the shrunken log_2_ fold change of total RNA.

### Principal component analysis

For total RNA, count data for genes was obtained as described in the method section on *Total RNA differential gene expression.* For labelled RNA, we used the output of GRAND-SLAM which reports the number of reads with at least 2 characteristic conversions as input for count data. Count data was transformed and normalised using in the same manner for total and labelled RNA. First count data was transformed and normalised with the variance-stabilising transformation (VST) implemented in DESeq2 (*62*). To focus our analysis on the most informative genes, we filtered for the top 500 genes with the highest variance before performing principal component analyses (PCA). PCA and gene loading plots, which illustrate the relative contribution of each gene to a given principal component, were generated from normalised count matrices.

### Statistical analysis

Western blots are analysed using BioRad Image Lab to acquire band intensities. Image J was used to acquire axonal density for regeneration experiments. For both, GraphPad Prism 6 software was used for all statistical analyses. One-way ANOVA was used, unless otherwise specified. All figure legends contain details of statistical tests and sample sizes. At least 2 different batches of differentiation were performed to acquire the i^3^ LMNs used in these experiments to control for batch-to-batch variability of neurons.

## DATA AND CODE AVAILABILITY

RNA sequencing data are available through the ArrayExpress under accession E-MTAB-14537. The mass spectrometry proteomics data have been deposited to the ProteomeXchange Consortium with the PRIDE, partner repository with the dataset identifier PXD046127. The codes used in this study are deposited in https://github.com/aleighbrown/lmn_bdnf. No new resources were generated in this study. BioRender was used to generate illustrations within figures.

## Acknowledgements

We thank Dr. Ling Hao for her guidance in mass-spectrometry analyses. We thank Professors Dario Alessi and Carsten Janke for their comments on the manuscript.

## Author Contributions

Conceptualization: JNSV, GS Data curation: ALB, SBS

Funding acquisition: JNSV, JNS, PF, GS Formal analysis: JNSV, ALB, KS, SBS

Investigation: JNSV, ALB, KS, CH, BG, DVC,SBS, ZM, AT, ML, LM, SM

Visualization: JNSV, ALB, KS, SBS Software: ALB, SBS

Supervision: JNSV, JNS, ALB, MS, AS, PF, GS

Writing – Original Draft: JNSV, ALB

Writing – Review & Editing: JNSV, ALB, KS, GS

---

**Figure S1.**
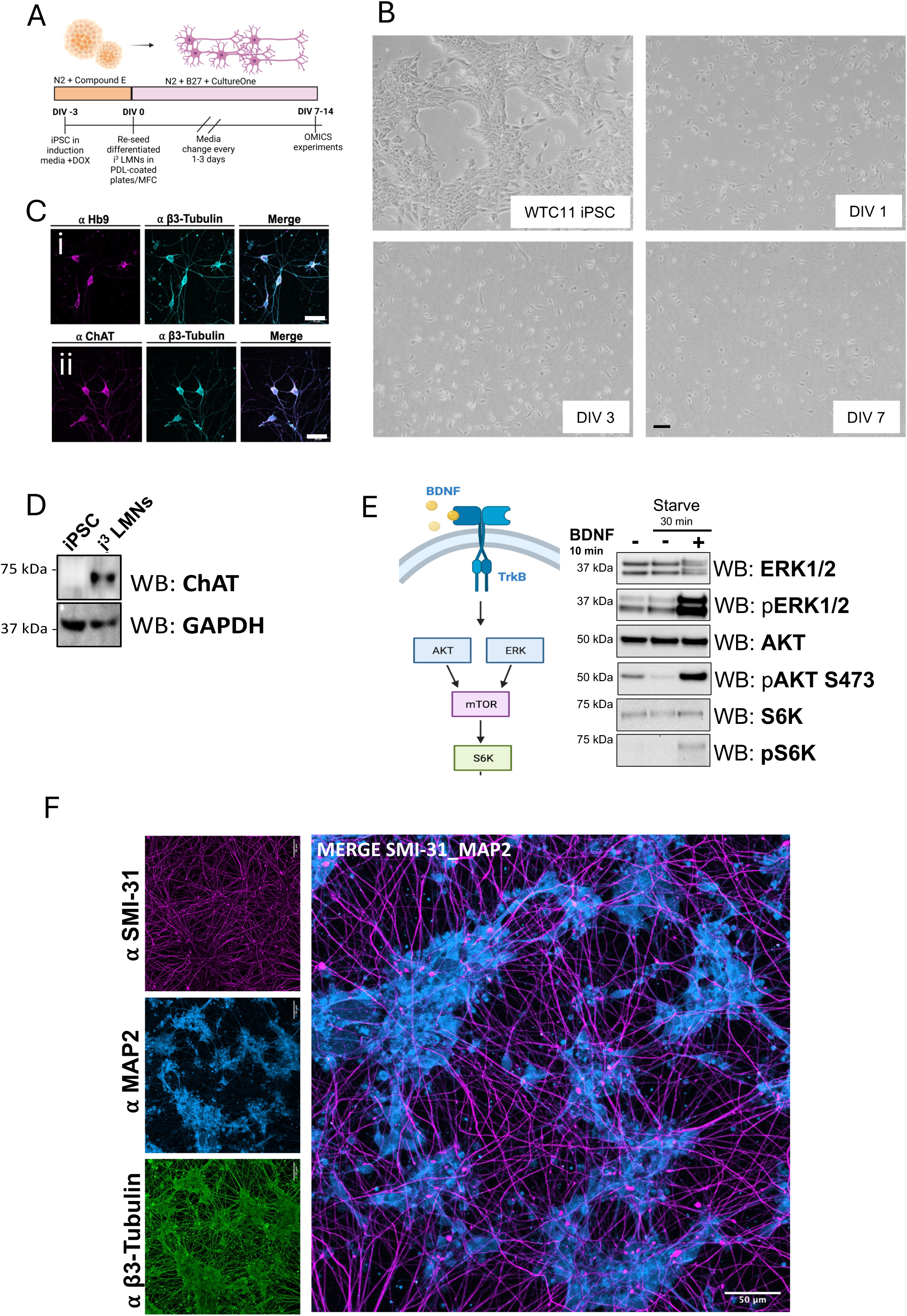
i3 LMNs express motor neuron markers and properly respond to BDNF treatment. A) Schematic of differentiation protocol of i^3^ LMNs. B) Brightfield images of WTC11 iPSC, and DIV1, 3, and 7 i^3^ LMNs post-differentiation. Scale bars: 50 μm. C) Immunofluorescence staining of i3 LMNs for motor neuron markers ChAT and Hb9. D) Western blot analysis of iPSC and differentiated i^3^ LMNs at DIV7 for expression of ChAT. E) Representation of BDNF signalling pathways (left). Western blot analysis of BDNF signalling in i^3^ LMNs after serum starvation for 30 min and BDNF treatment (25 ng/mL) for 10 min. F) Immunofluorescence of i^3^ LMNs (DIV 12) cultured in tripartite MFCs. SMI-31 signal delineates the axonal compartment and MAP2 labels the somatodendritic compartment.

**Figure S2.**
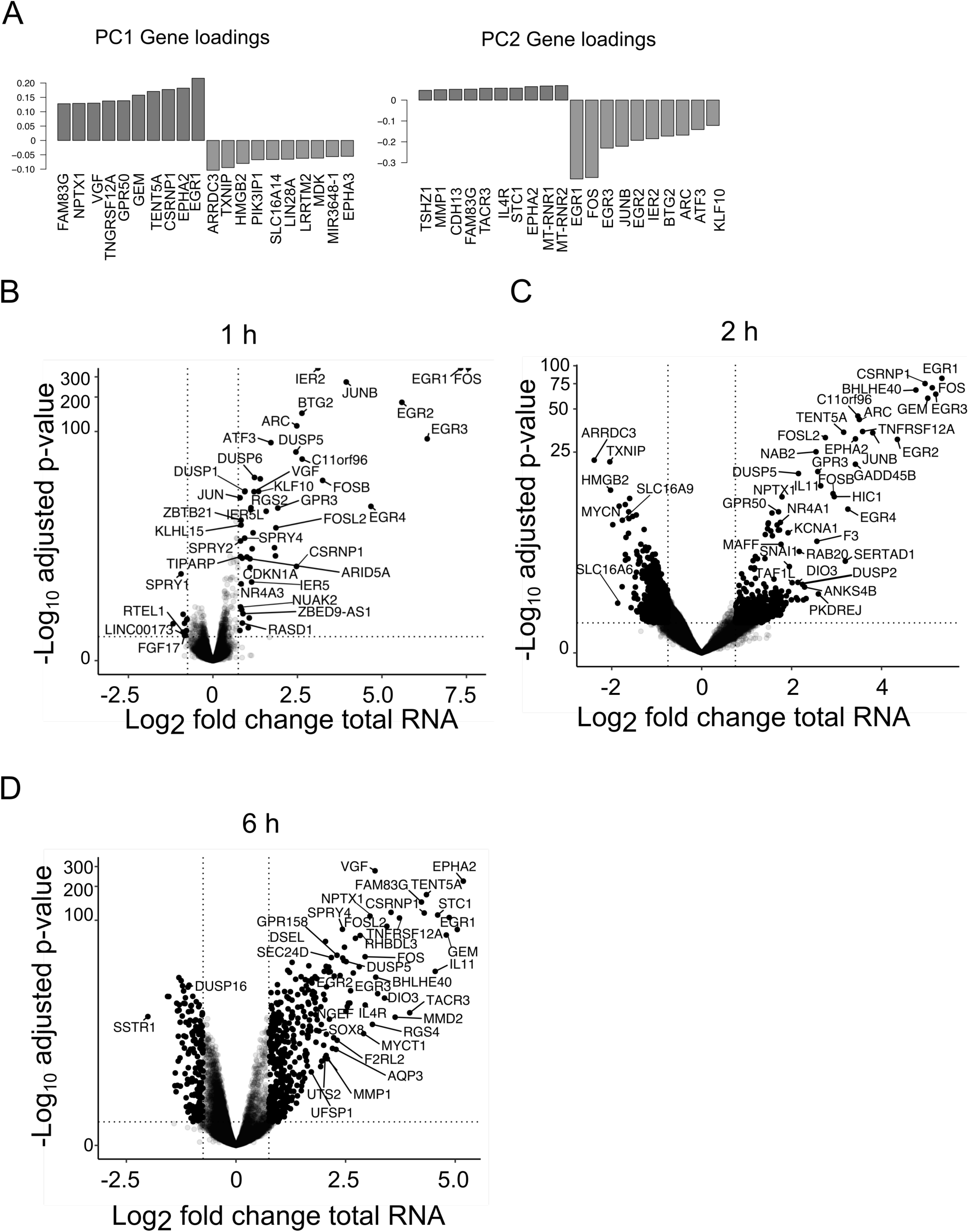
BDNF differentially regulated genes in human motor neurons. A) Gene loading plots for Fig. 1B showing the most important genes for the first 2 principal components. Larger loading values represent a greater contribution to principal component. B-D) Volcano plots of differentially expressed genes within 1 h (B) (58 DE genes, 48 upregulated, 10 downregulated), 2 h (C) (2,131 DE genes, 1,172 upregulated, 959 downregulated), and 6 h (D) (604 DE genes, 434 upregulated, 170 downregulated) of BDNF treatment in i3 LMN. Lines define cutoff for a gene being called as differentially expressed (adjusted p-value < 0.1, abs (log2 fold change) > 0.75). Y-axis is cut for genes with p-values approaching zero. N=3 biological replicates per condition and labelling time.

**Figure S4.**
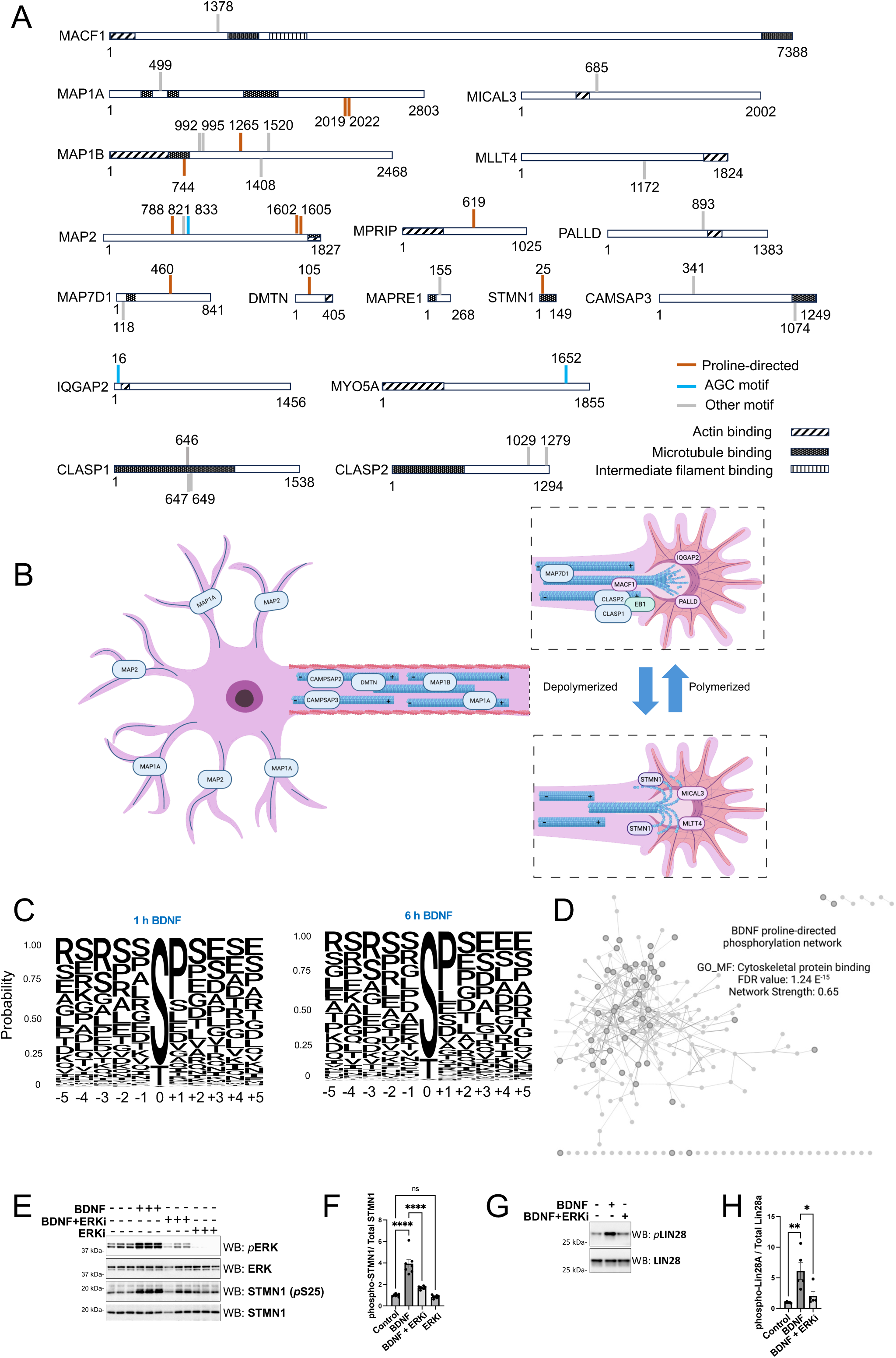
Phosphorylation patterns of cytoskeletal-binding proteins by BDNF. A) Significant phosphorylation events on proline-directed (pS/TP) and AGC (RXRXXpS/T) motifs downstream of BDNF treatment on microtubule and actin-binding proteins. B) Schematic of cytoskeletal-binding proteins identified as BDNF phospho-substrates in the context of neuronal localization/enrichment. C) Sequence logo of differentially phosphorylated sites after 1 h (left) and 6 h (right) of BDNF treatment. The height of letters displays the frequency of the amino acid at each site. D) Cytoscape StringdB network of BDNF-dependent proline-directed serine residue enrichment for cytoskeletal protein-binding (in yellow). Inputs peptides p-value < 0.05. E) Western blot analysis of phosphorylated STMN1 (S25) in DIV7 i^3^ LMNs after 25 ng/ml BDNF with or without the ERK inhibitor U0126 (20 μM) for 1 h. F) Quantification of the western blot analysis shown in (E); N=7. G) Western blot analysis of phosphorylated LIN28A (S200) after 25 ng/mL BDNF with or without the ERK inhibitor U0126 (20 μM) for 1 h. H) Quantification of the western blot analysis shown in (G); N=5. Western blot analyses were performed with neurons derived from at least two independent differentiations. One-way ANOVA. Error bars: SEM. p value: ∗ = < 0.05; ∗∗ < 0.01; ∗∗∗ < 0.001; ∗∗∗∗ < 0.00001.

**Figure S5.**
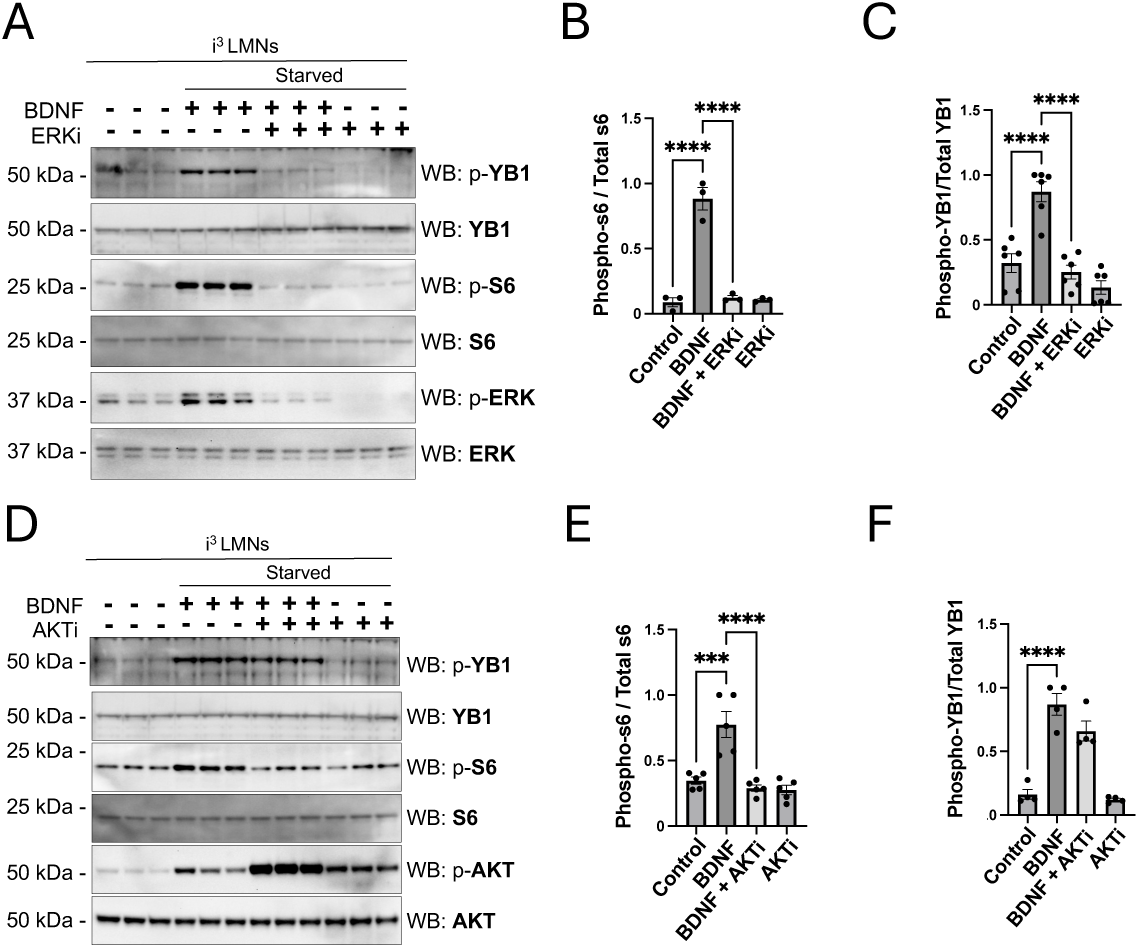
BDNF activation of RSK and S6K signalling in human motor neurons. A) Western blot analysis of RSK and S6K kinase signalling downstream of the BDNF-ERK pathway. i^3^ LMNs were serum starved for 2 h before BDNF treatment (1 h; 25 ng/mL), with or without co-treatment of ERKi (U0126; 20 μM). B) Quantification of the western blot analysis for phospho-YB1 shown in (A). N=6. C) Quantification of the western blot analysis for phospho-s6 shown in (A). N=6. D) Western blot analysis of RSK and S6K kinase signalling downstream of the BDNF-AKT pathway. i^3^ LMNs were serum starved for 2 h before BDNF treatment (1 h; 25 ng/mL), with or without co-treatment of ERKi (Ipatasertib; 1 μM). E) Quantification of the western blot analysis for phospho-YB1 shown in (D). N=5. F) Quantification of the western blot analysis for phospho-s6 shown in (D). N=5. All western blot analyses were performed with neurons derived from at least two independent differentiations. One-way ANOVA. Error bars: SEM. p value: ∗ = < 0.05; ∗∗ < 0.01; ∗∗∗ < 0.001; ∗∗∗∗ < 0.00001.

**Figure S6.**
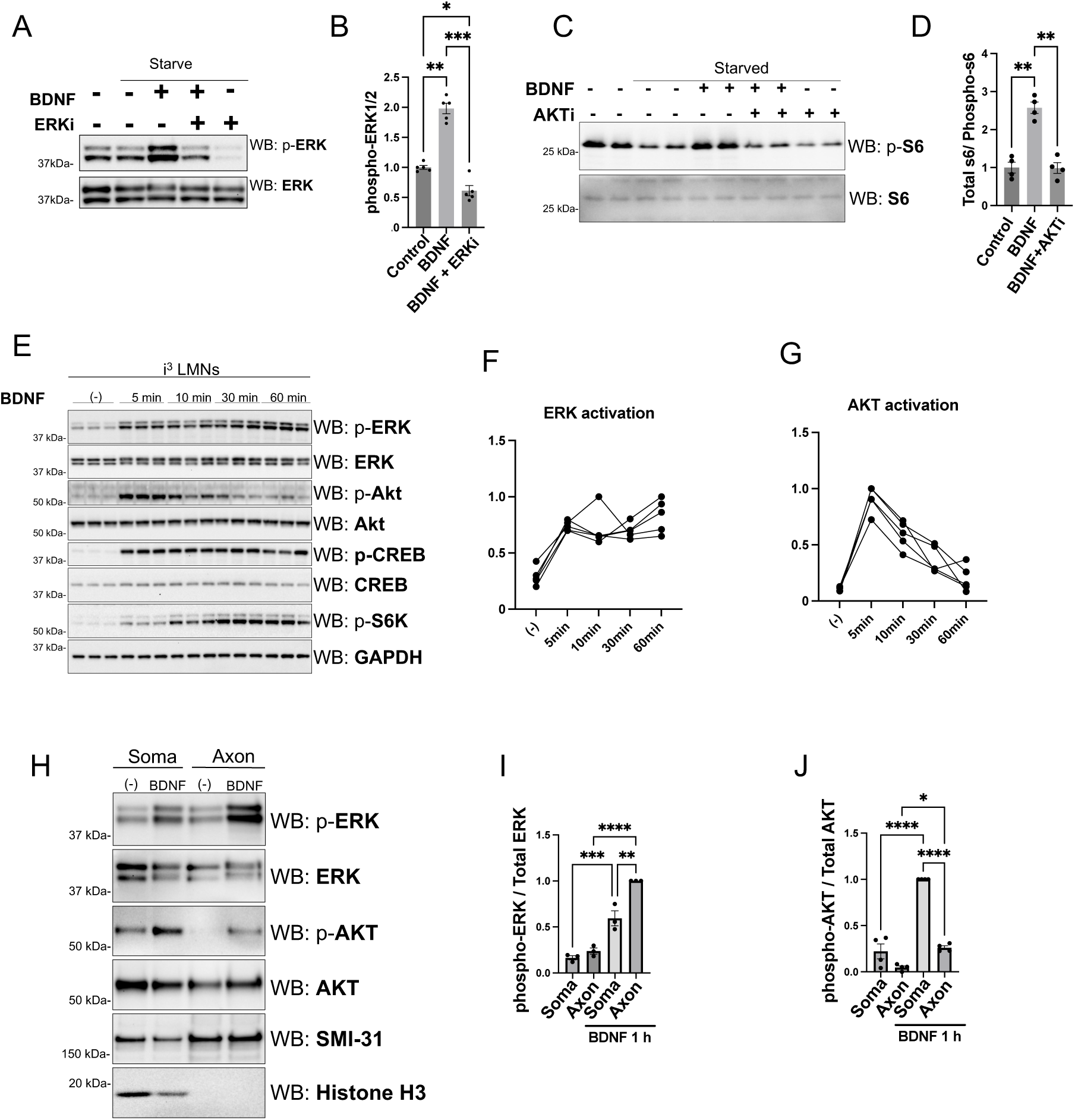
ERK and AKT signalling in axons of BDNF-treated motor neurons. A) Western blot analysis of phospho-ERK1/2 after treatment of BDNF (25 ng/mL) and co-treatment with ERKi (U0126 20 μM). B) Quantification of the western blot analysis in (A) N=5 C) Western blot analysis of phospho-S6K after treatment of BDNF (25 ng/mL) and co-treatment with AKTi (Ipatasertib 0.1 μM). D) Quantification of the western blot analysis shown in (C). N=5. E) Western blot analysis of BDNF (25 ng/mL) signalling kinetics. F) Quantification of ERK activation across timepoints shown in (E). Each dot represents an independent experiment. N=5 per time point. G) Quantification of AKT activation across timepoints shown in (E). Each dot represents an an independent experiment. N=5 per time point. H) Western blot analysis of somatic and axonal proteins isolated from i3 LMNs cultured in MFCs. SMI-34 and histone H3 were used as markers for axonal and somatic enrichment, respectively. BDNF (1 h; 25 ng/mL). I) Quantification of somatic and axonal activation of ERK1/2 by BDNF from (H). J) Quantification of somatic and axonal activation of AKT by BDNF from (H). All western blot analyses are performed with neurons derived from at least two differentiation rounds. One-way ANOVA. Error bars: SEM. p value: ∗ = < 0.05; ∗∗ < 0.01; ∗∗∗ < 0.001; ∗∗∗∗ < 0.00001; ns., not significant.

